# Functional traits explain growth resistance to successive hotter droughts across a wide set of common and future tree species in Europe

**DOI:** 10.1101/2024.07.04.602057

**Authors:** Lena Kretz, Florian Schnabel, Ronny Richter, Anika Raabgrund, Jens Kattge, Karl Andraczek, Anja Kahl, Tom Künne, Christian Wirth

**Author notes:** LK and FS contributed equally and shared first authorship.

## Abstract

- In many regions worldwide, forests suffer from climate change-induced droughts. The ‘hotter drought’ in Europe in 2018 with the consecutive drought years 2019 and 2020 caused large-scale growth declines and forest dieback. We investigated if tree growth responses to the 2018–2020 drought can be explained by tree functional traits related to drought tolerance, growth, and resource acquisition.
- We assessed growth resistance, that is, growth during drought compared to pre-drought-conditions, in 71 planted tree species using branch shoot increments. We leveraged gap-filled trait data related to drought tolerance (P50, stomata density and conductivity), growth and resource acquisition (SLA, LNC, C:N, A_max_) and wood density from the TRY database to explain growth resistance for gymnosperms and angiosperms.
- We found significantly reduced growth during drought across all species. Legacy effects further decreased growth resistance in 2019 and 2020. Gymnosperms showed decreasing growth resistance with increasing P50 and acquisitiveness, such as high SLA, LNC, and A_max_. Similar results were found for angiosperms, however, with less clear pattern. Four distinct response types emerged: ‘Sufferer’, ‘Late sufferer’, ‘Recoverer’ and ‘Resisters’, with gymnosperms predominately falling into the ‘Sufferer’ and ‘Late sufferer’ categories.
- Our study provides evidence for significant growth reductions and legacy effects in response to consecutive hotter droughts, which can be explained by functional traits across a wide set of tree species. The a posteriori classification into response types revealed the diversity of temporal responses to a prolonged drought. We conclude that high drought tolerance bolsters growth resistance, while acquisitive species suffer stronger under drought.

## Introduction

In many regions of the world forest productivity decreases, as trees suffer from more intense and frequent drought events caused by climate change (Allen et al., 2010; IPCC, 2014; McDowell et al., 2020). Negative impacts on forests are particularly pronounced for so-called ‘hotter droughts’ which are compound events characterized by low precipitation and simultaneous heat waves (Allen et al., 2015). Such hotter droughts cause enhanced soil-water depletion and increases in canopy temperature, potentially surpassing physiological tolerance thresholds and thereby inducing strong growth reductions (Allen et al., 2015; Buras, Rammig, & Zang, 2020). Such growth declines often precede large-scale tree mortality events, eventually being amplified by climate change-induced insect and pathogen outbreaks. (Allen et al., 2015; McDowell et al., 2020). Thus, hotter droughts negatively impact many ecosystem functions and services of forests, such as carbon sequestration (Buras, Rammig, & Zang, 2020; Senf et al., 2020) and transpirative cooling (Richter et al., 2021), and may induce strong changes in species compositions (Schuldt et al., 2020). However, we thus far have only a limited understanding of how intense drought events cause growth reductions across a wide range of tree species. We further lack knowledge on functional properties of tree species which help to develop trait-based models to generalize responses to hotter drought across wide taxonomic gradients and to parametrize models predicting growth responses and eventually mortality risks (Adams et al., 2017).

In the year 2018, Central Europe experienced a hotter drought, which was climatically the most extreme drought since the beginning of climatic records in Europe (Schuldt et al., 2020; Zscheischler & Fischer, 2020). The hotter drought conditions persisted in the year 2019 and in many Central European regions continued even until 2020 (Rakovec et al., 2022). These three consecutive drought years (hereafter referred to as the ‘2018–2020 drought’) may mark the beginning of a new era of compound climate extremes which is in line with models of climate change that project hotter, drier and more extreme climatic conditions, particularly in summer months, for Central Europe during the 21^st^ century (IPCC, 2014; Reichstein et al., 2013; Samaniego et al., 2018; Trenberth et al., 2014; Zscheischler & Seneviratne, 2017). Such consecutive and hotter droughts induce prolonged stress, amplified reductions in tree growth and eventually large-scale forest dieback, especially when interacting with fungal pathogen and insect outbreaks (Hari et al., 2020; Kleine et al., 2021; Schnabel et al., 2022; Thonfeld et al., 2022).

Here we aim to understand tree growth responses across a broad range of native and introduced Central European tree species to the 2018–2020 drought. We included introduced species from North America and Asia, based on their current relevance in Central Europe as future tree species under climate change. Growth resistance in this context is defined as the ratio of the growth during the drought years and the growth prior to the drought. Especially interesting are potential growth reactions during the second and third consecutive drought year (2019 and 2020), as droughts can still affect trees negatively one to five years after the actual drought event, which is known as drought legacy effect (Anderegg et al., 2015; Anderegg, Kane, et al., 2013; Bigler et al., 2006; Gazol et al., 2020; Kannenberg et al., 2018; Schnabel et al., 2022). Legacy effects of the 2018 drought, such as associated damages to the water transport system of trees (Anderegg, Plavcová, et al., 2013), may limit their capacity to deal with and recover from the subsequent drought years. Moreover, growth reductions may be amplified by a cumulative build-up of soil water deficits. Hence, one may expect a lower growth resistance in the consecutive drought years 2019 and 2020, which is, for instance, consistent with recent reports of a lower growth resistance in 2019 compared to 2018 in a Central European floodplain forest (Schnabel et al., 2022). However, observational forest studies are typically restricted to relatively few tree species making it complicated to test for the trait-based mechanisms driving legacy effects and to generalize these across regional tree floras.

The recent years have seen a surge of studies exploring the trait-based mechanisms underpinning drought effects on tree growth (Bose et al., 2020; Larysch et al., 2022; Liu et al., 2022). These studies are typically restricted to few tree species in single sites or combine observations from different sites that vary in environmental conditions. Under water shortage, plants are facing a trade-off between carbon gain and water loss (Cowan & Farquhar, 1977). Thus, the physiological key processes that cause growth reductions are either carbon starvation or partial hydraulic failure, but the relative balance between both processes varies strongly between tree species and with growing conditions (Adams et al., 2017; McDowell et al., 2008; Sala et al., 2010; Schuldt et al., 2020; Sevanto et al., 2014). It emphasizes the importance to observe growth responses under drought in a single site under comparable conditions. Traits related to drought tolerance, such as P50 (pressure, where 50 % of the hydraulic system’s conductivity has been lost (Adams et al., 2017; Guillemot et al., 2022)), or stomatal control traits, may help to understand growth reductions caused by those two mechanisms (Schnabel et al., 2021, 2022). One may expect that tree species whose functional traits indicate a high drought tolerance, such as a low P50 indicating a high tolerance to negative water potentials (Jarbeau et al., 1995; Choat et al., 2018), show a higher growth resistance. Moreover, high stomata density can be caused by lower stomata size, but also may be related to specific spatial distribution (Klein, 2014; Lawson & Blatt, 2014), both possibly indicating a faster or more precise stomata control and, thus a better adaptation to drought under drought conditions. The stomata control is also expected to link to different adaptation strategies under drought of anisohydric and isohydric species (Klein, 2014; N. McDowell et al., 2008). Traits related to stomatal control are complex and depend on tree hydraulics, such as xylem and leaf water potential, but also on the photosynthetic rate. Isohydric species, for example, close their stomata earlier i.e., at lower water potentials or water pressure deficit, often have reduced mean stomata conductance to avoid hydraulic failure during drought and thus are considered water-savers. In contrast, as anisohydric species close their stomata late and thus often have higher mean stomata conductance, they are considered water-spenders (Klein, 2014; N. McDowell et al., 2008). Along this gradient of stomatal behaviour, we would expect that species with lower stomatal conductance are less susceptible to drought. Next to drought-tolerance traits, growth and resource acquisition related traits, such as traits of the leaf economics spectrum representing the slow-fast gradient of plant growth (Guillemot et al., 2022; Reich, 2014) may explain growth resistance to drought. First, tree species with LES trait expressions of the leaf economic spectrum (LES, Díaz et al., 2016) related to conservative resource use and slow growth, such as a high carbon to nitrogen ration (C:N), may feature a higher growth resistance to drought (Choat et al., 2015; Reich, 2014; Wright et al., 2004). This view is consistent with reports of a high correlation of these traits with traits related to cavitation resistance such as P50 (Guillemot et al., 2022; Reich, 2014; Schnabel et al., 2021). In contrast, LES traits related to acquisitive resource use and fast growth, such as high specific leaf area (SLA), leaf nitrogen content (LNC), and light-saturated maximum photosynthetic rate (A_max_), may feature a lower growth resistance to drought (Wright et al., 2004; Reich, 2014; Díaz et al., 2016; Greenwood et al., 2017). In addition, wood density combines various wood properties and is associated with mechanical strength and water transport of the stem (Chave et al., 2009; Zanne et al., 2010). Divergent effects were found before. While some found that species with high wood density have lower mortality rates during drought (Greenwood et al., 2017) and higher growth resistance (Serra-Maluquer et al., 2022), others found for temperate angiosperms higher canopy dieback with high wood density (Hoffmann et al., 2011). Still, slow-growing species tend to have denser wood (Chave et al., 2009; L. Poorter, 2008), thus we would expect growth resistance to increase along with wood density.

To guide management decisions and to improve the predictive capacities of forest models it is important to understand the response of all Central European tree species to the 2018–2020 drought, i.e. not only of those dominating today, but also the many subordinate or biogeographically neighboring tree species that may form the forest under future climate regimes. Currently, establishing the relationship between functional properties of tree species and their responses to the novel climate situation is challenging. To gain enhanced understanding, we have to exploit the unique sequence of climate events since 2018 and find means to reconstruct tree responses for as many tree species with relevance for Central Europe, including the native tree flora and common non-native tree species. For this purpose, national forest inventories are of limited use for two reasons: (i) they do not possess the necessary temporal resolution to capture the sequence of growth responses (initial resistance, legacy effects, potential recovery), (ii) the Central European managed forest landscape is dominated by few merchantable tree species such as Norway Spruce (*Picea abies*), Scots Pine (*Pinus sylvestris*), European Beech (*Fagus sylvatica*) and Pedunculate Oak (*Quercus robur*), which make up 73.5% of the forest area according to the last German national forest inventory (BWI 2012). In contrast, rare species with large potential for forestry under drier and hotter climates, such as the Checker Tree (*Sorbus torminalis*) and Downy Oak (*Quercus pubescens*) are hardly captured (Buras & Menzel, 2019; Kunz et al., 2018). For instance, three species, as reported by Schnabel et al. (2022), showed reduced growth resistance and drought legacy effects in 2019 compared to 2018, but such observational studies are typically restricted to few tree species making it difficult to derive generalizable conclusions on the trait-based mechanisms across tree species which may explain this drought legacy effect.

Here, we examine the effects of the three consecutive drought years 2018–2020 on a large set of 71 planted tree species (Table S1) under experimental conditions in the research arboretum ARBOfun. The arboretum contains 100 species including gymnosperms and angiosperms as well as native and common exotic species. Each species is 5 times replicated in a wide stand with no competition and grown under similar soil conditions. ARBOfun was designed to study responses to climate variability for a large number of tree species. Taking advantage of this unique design, we here aim to provide new insights into the growth resistance of an unprecedented set of tree species and to test if the strength and type of growth responses can be predicted by tree functional traits related to drought tolerance and resource acquisition capacity. We hypothesized that:

1. The 2018-2020 drought reduced tree growth, with a greater reduction in growth resistance in the years 2019 and 2020 due to legacy effects.
2. Tree species whose functional traits indicate drought tolerance show a higher growth resistance to drought stress than drought intolerant species.
3. Tree species whose resource acquisition traits favour rapid growth are more susceptible to drought and show a lower growth resistance during drought than tree species with traits indicating a conservative resource use.

## Material and Methods

### Experimental design and study site

The ARBOfun research arboretum is located south of Leipzig (Saxony, Germany, 51°161N, 12°301E). The experiment was established in 2012 on 2.5 ha of former extensively used arable land with the soil type Luvisol. In 2012 a set of 69 species were planted, and 31 additional species were added in 2014, totalling 100 tree species. Each species is randomly replicated 5 times within a block design, where each block contains one individual per species (Figure 1). The tree individuals are arranged in a checkerboard-pattern with a wide spacing 5.8 m to prevent competition in the early years of the experiment. Due to mortality, predominantly unrelated to drought (e.g. vole damage to roots), not all species have five replicates. The meadow between the trees is mown twice per year. The selected tree species represent the diversity of woody species native to Europe, originating from the gradient from hemi-boreal to sub-mediterranean forests and, in addition, includes selected species from North-America and Asia frequently planted in forest plantations or cities (Table S1).

**Figure 1:**
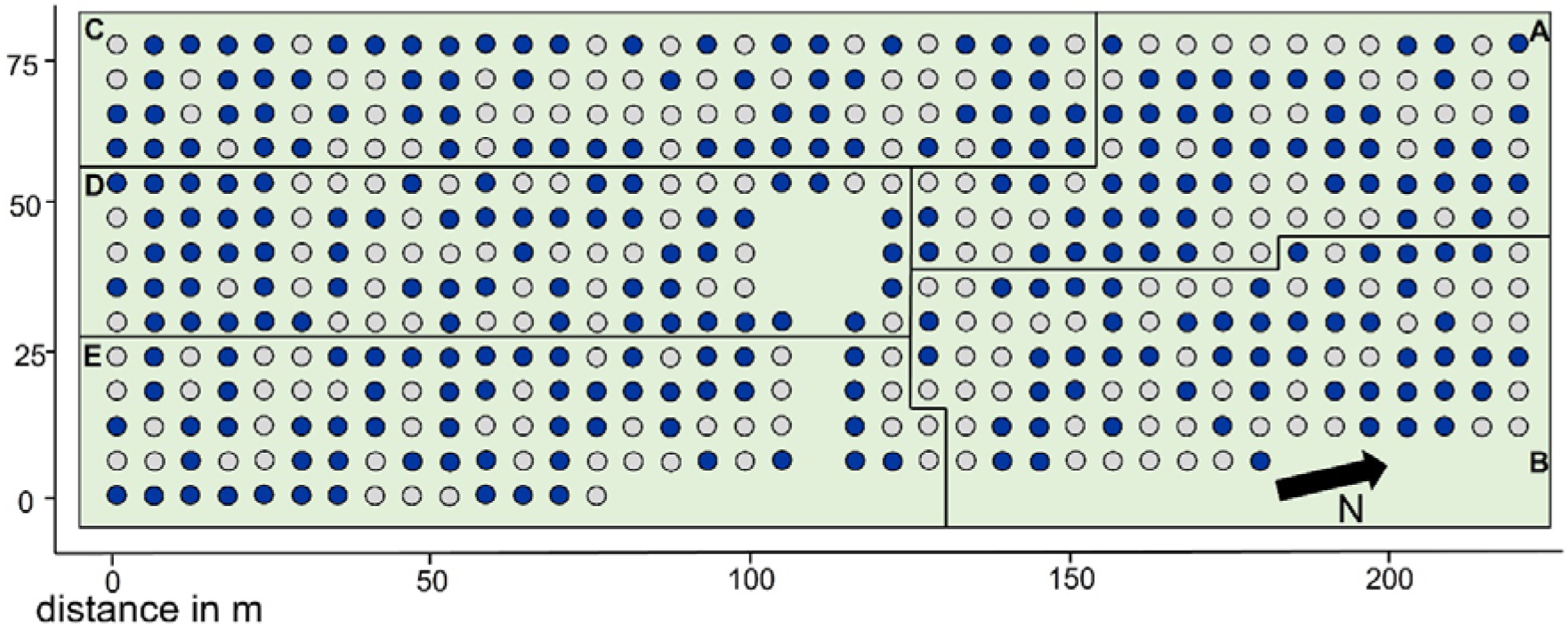
Study site. Top view on the ARBOfun study site. The points represent the 100 species, each randomized within 5 blocks (A-D). The dark blue colour marks the trees used for this study. The planting distance between trees is 5.8 m.

The study site is located at an elevation of 150 m a.s.l. in the transition zone from maritime to continental climate. At the area the mean annual precipitation is approximately 520 mm, and mean annual temperature is 9.7 °C (1980-2020; DWD Climate Data Center [CDC], Station Leipzig/Halle, ID 2932). In 2018–2020 a period of consecutive drought and heat occurred all over Central Europe. To characterise the climatic conditions at our study site, we examined monthly temperature and precipitation as well as the standardized water balance of precipitation minus potential evapotranspiration using the Standardized Precipitation Evapotranspiration Index (SPEI; Vicente-Serrano, et al. 2010). SPEI is an often-used drought index (Hari et al., 2020; Schwarz et al., 2020) which quantifies drought severity according to a drought’s intensity and duration across time scales (Vicente-Serrano et al., 2010). We calculated three different SPEI lengths with the SPEI package (Beguería & Vicente-Serrano, 2017) in R: SPEI3 capturing the water balance during the main vegetation period (Mai–July), SPEI6 during the full vegetation period (April–September) and SPEI12 during the entire year (January–December). Monthly climate data were derived from the weather station located closest to the experiment that featured complete records (DWD Climate Data Center [CDC], Station Leipzig/Halle, ID 2932). Potential evapotranspiration was calculated with the FAO-56 Penman-Monteith equation (Beguería & Vicente-Serrano, 2017) using the following DWD data: monthly means of daily minimum temperature, daily maximum temperature, wind speed, cloud cover, atmospheric surface pressure, relative humidity, vapor pressure as well as station elevation and latitude.

### Tree sampling and trait measurements

For the present study, we measured shoot increments for a total set of 71 tree species (Table S1). The measurements took place in spring 2021. We used the scars of bud scales to retrospectively measure the shoot increments of three lateral branches per tree from the year 2020 back to the year 2016 (Figure S1). For the measurement of the lateral branches, first, the lateral branch that was south-facing and at about ¼ total tree height was selected and measured. Additionally, a second lateral branch was selected anti-clockwise around the tree 120° angle from the first branch, while a third lateral branch was selected in the same way starting from the second branch. For the present study, we included only species with at least 2 replicates each with a minimum of 2 branch measurements which could at least be dated back until the year 2017. This leaves us with a total of 850 measured branches on 284 tree individuals.

Species resistance is defined as the lack of an ecological performance reduction during disturbance or stress conditions (Kaufman, 1982; MacGillivray & Grime, 1995). We used resistance as indicator for drought stress of the trees and calculated it as the ratio of performance during the disturbance/stress and before the disturbance/stress according to (Lloret et al., 2011).

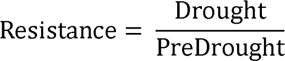

‘Drought’ corresponds to growth during one or all of the drought years 2018, 2019, or 2020, while ‘PreDrought’ correspond to the growth before the drought, which we calculated as the mean of the reference years 2016 and 2017 (growth for 2016 was only available for 81 % of the branches). We calculated growth resistance as:

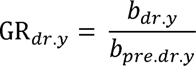

Where, GR is the growth resistance, *b* the length of the branch increment, while *dr.y* is the drought year and *pre.dr.y* the pre-drought reference. Note, that values of GR*_dr.y_* > 1, also indicate resistance even though the growth rates exceed the pre-drought performance.

We selected eight different functional traits that we, based on the literature (Anderegg et al., 2019), expected to be key traits for growth resistance and that are available in the TRY database for a large number of species (Kattge et al., 2020): (1) P50, (2) stomata density, (3) stomatal conductance, (4) specific leaf area (SLA), (5) leaf nitrogen content (LNC), (6), leaf carbon to nitrogen ratio (C:N), (7) maximal photosynthetic rate (A_max_), and (8) wood density.

1. P50 (MPa) describes the xylem pressure, where 50 % of the hydraulic system’s conductivity has been lost. If the xylem pressure falls below that, the plant is exposed to a high risk of lethal embolism (Brodribb & Cochard, 2009; Sperry & Tyree, 1988).
2. Stomata density (mm^−2^) is the number of stomata per leaf area and can be linked to stomata size and distribution, but also indicates stomatal control and conductance (Klein, 2014).
3. Mean stomatal conductance (mmol m^−2^ s^−1^) is the conductivity for water vapor per leaf area per time of the stomata and can be linked to tree hydraulics and leaf water potential, but also to the photosynthetic rate and with that mechanistically to an acquisition strategies (Garcia-Forner et al., 2016).
4. Specific leaf area (SLA, mm^2^ mg^−1^) is leaf area gain per invested leaf biomass. It is suggested to be negatively related to plant performance under drought (H. Poorter et al., 2009). Further it is a key trait representing resource acquisition and a fast growth and resource acquisition of the plant economic spectrum (Reich, 2014; Wright et al., 2004).
5. Leaf nitrogen content (LNC, mg g^−1^) is a major component of photosynthetic compounds such as the enzyme Rubisco and thus directly affects the photosynthetic capacity of leaves (Evans, 1989; Reich et al., 1995) and is also one of the traits representing a fast growth and resource acquisition of the plant economic spectrum (Reich, 2014; Wright et al., 2004).
6. The leaf carbon to nitrogen ratio (C:N) is, beside others functions, linked to growth, but also stress responses (Hessen et al., 2004). A high C:N indicates a low N concentration, thus slow growth as mentioned before, but also a high C concentration, which can also indicate thicker cell wall, which makes the species more resistant to drought stress (Reich, 2014; Wright et al., 2004).
7. Light-saturated maximum photosynthetic rate (A_max_, μmol g^−1^ s^−1^) is the maximum carbon assimilation rate under normal water conditions as an index of photosynthetic capacity (Anderegg et al., 2018; Zhu et al., 2018), and associated with fast resource acquisition (Lambers & Poorter, 2004).
8. Wood density (g cm^−3^) combines diverse wood properties, such as mechanical strength, water storage and transport (Chave et al., 2009; Zanne et al., 2010). In general, the wood density strongly depends on the porosity group, however, we would expect an overall trend that species with high wood density are more resistant against drought.

### Statistical analysis

All statistical analysis were done with the statistical software R (R Core Team, 2020). We used linear mixed-effects models (lme function in nlme package, (Pinheiro et al., 2023) to predict growth resistance across tree species. We used drought year (2018, 2019 and 2020) coded as factor as a fixed effect. We log-transformed tree growth resistance to fulfil model assumptions (normality and homogeneity of variance) and used branch ID nested within tree ID as a nested random effect structure. As reference of tree growth under ‘normal’ climatic conditions, we used the mean growth in 2016 and 2017 (which were neither exceptionally wet nor dry years, nor were the years before, which could have caused legacy effects in 2016 and 2017). We used a post-hoc test for comparisons between the years (emmeans function in the emmeans R package, (Lenth, 2023), corrected for multiple comparisons with first order autocorrelation structure (corAR1) with the year as covariate. We also ran linear mixed-effects models in the same way for gymnosperms and angiosperms separately and for every single species.

We used available trait data from the TRY Plant Trait Database (Kattge et al., 2020). Since the available data do not compile a complete dataset, we conducted a gap-filling to predict trait values for those traits and species that were not available. For the gap-filling we used a hierarchical Bayesian implementation of probabilistic matrix factorization (BHPMF, Schrodt et al., 2015). In a first step we cleaned the available data on TRY, excluded non-vascular species, juveniles, and non-healthy plants. We further excluded outlier values with a distance of > 5 standard deviations from taxonomic or functional group means (Kattge et al., 2011, 2020) and A_max_ and stomatal conductance measured under conditions of CO_2_ not ambient (300-450 ppm), unsaturated light conditions (< 800 µmol m^−2^ s^−1^) and temperature outside 20 and 30 °C. Then we z-transformed the data and ran the gap-filling with the BHPMF approach. In a post-processing we back-transformed the data and excluded data of the 25% quantile with highest standard deviation per prediction of a trait record (Fazayeli et al., 2014). We excluded data with a distance of > 3 standard deviations from taxonomic and functional group means.

With the gap-filled data we ran linear mixed-effects models (lme function in nlme package (Pinheiro et al., 2023), to predict growth resistance by each individual trait interaction with drought year (2018, 2019 and 2020, coded as factor). We again used the branch nested in the tree as random factor and the mean growth in 2016 and 2017 as a reference. We also ran principal component analyses (PCA). For the PCAs we could include 53 species with full trait coverage, 18 gymnosperms and 35 angiosperms. The PCAs where conducted with the prcomp function. We also used the loadings of PCA axes 1 and 2 as predictors in the same way to predict growth resistance with linear mixed effect models. We used Fisher’s exact test for contingency table data for analysing different response types within the clades.

## Results

### Climate

We observed consecutive hotter drought conditions from 2018-2020, with climatic drought severity declining slightly from 2018 over 2019 to 2020 (SPEI12 values of −2.06, −1.76, −1.53, respectively, Figure 2A). All three years were among the driest in the last 40 years when considering the peak vegetation period (May-July), the full vegetation period (April-September), and the entire year (Figure 2 and Figure S2)). However, especially the coincidence of high temperatures (Figure 2B) and low precipitation (Figure 2C) as well as the consecutive nature of these droughts marked the 2018-2020 drought as exceptional.

**Figure 2:**
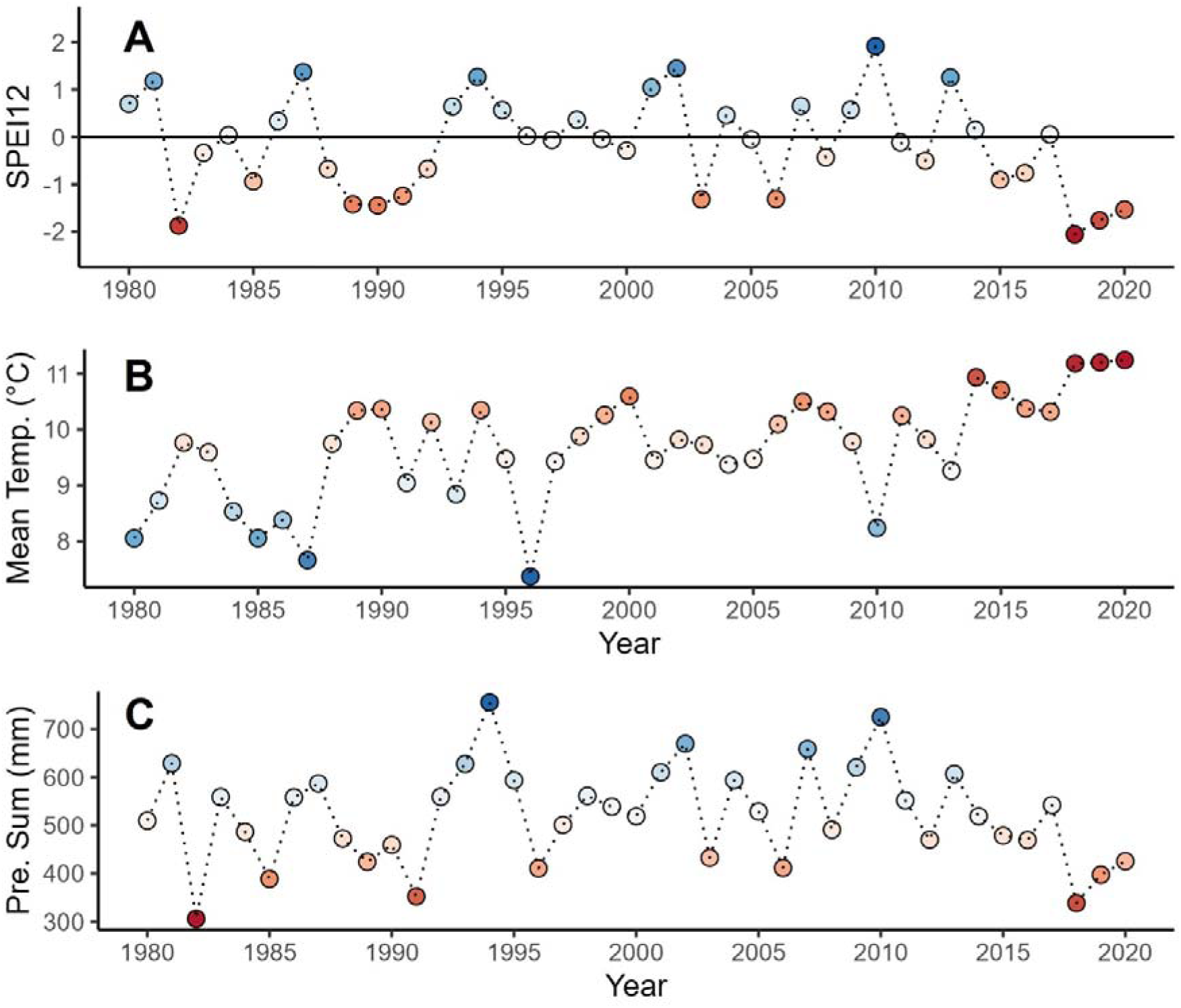
Standardised Precipitation-Evapotranspiration Index (SPEI), mean annual temperature and annual precipitation sum. The SPEI12 is calculated for the whole year from January to December. The zero line is the reference period 1981–2010. Blue coloured dots indicate positive SPEI values, while red coloured dots show negative SPEI values. The mean temperature and precipitation sum are calculated over the whole year. Also, here, the blue and red colour indicates higher and lower values compared to the reference period, respectively.

### Growth resistance of all species

The lowest growth resistance of 0.018 was measured for *Crataegus monogyna* in 2020, while the highest growth resistance of 6.962 was measured for *Fraxinus excelsior* also in 2020. Median growth resistance per species varied from 0.318 (*Juglans regia*) up to 1.314 (*Castanea sativa*, Figure 3), while the gymnosperm with lowest median growth resistance of 0.436 was *Tsuga canadensis* and with highest median growth resistance of 1.143 was *Pinus mugo*.

**Figure 3:**
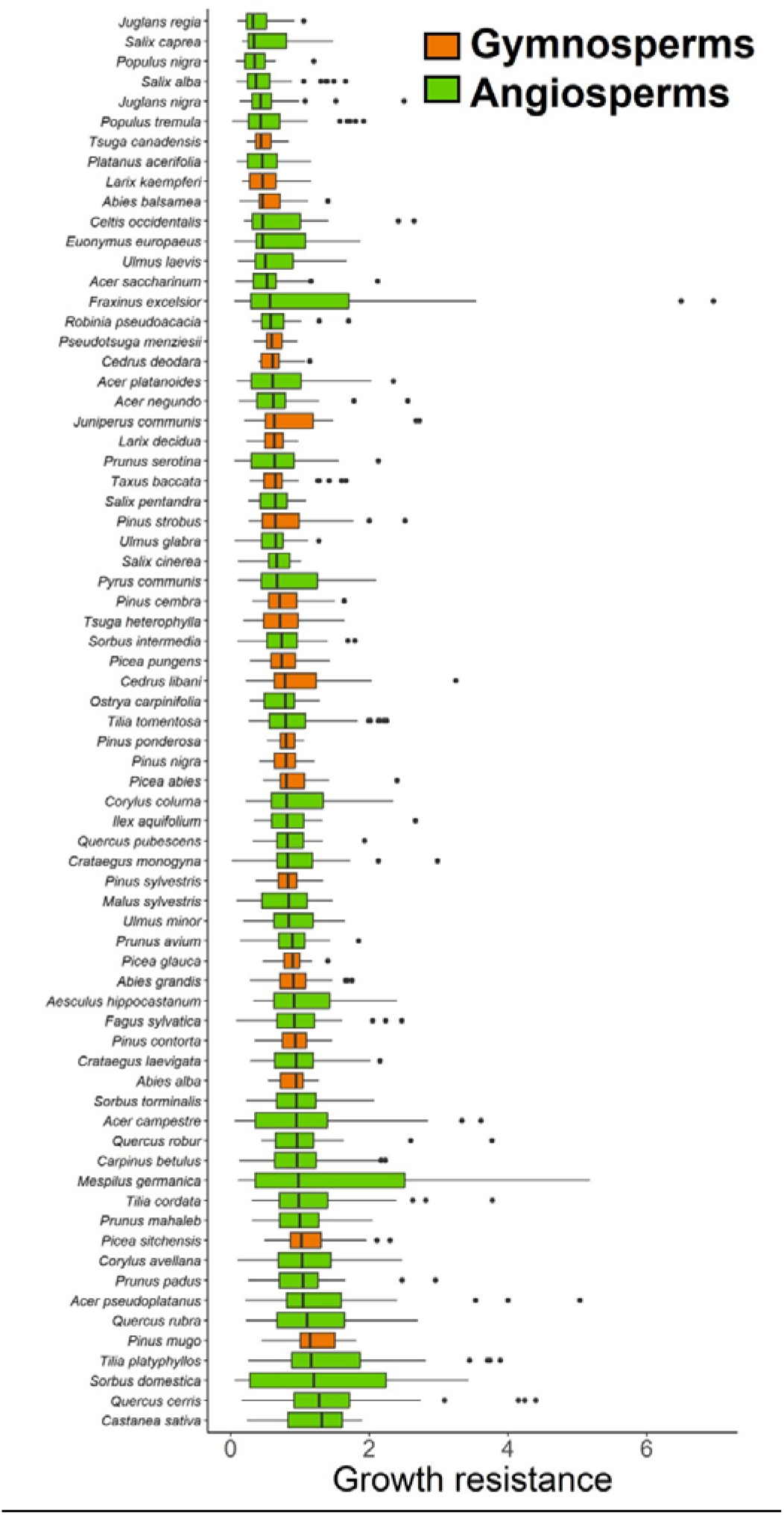
Growth resistance of all tree species, sorted by the median growth resistance per species.

We found that the growth resistance across all species was significantly reduced in the years 2018 (*p*<0.001), 2019 (*p*<0.001), and 2020 (*p*<0.001) compared to the mean growth of the reference years 2016 and 2017 (Figure 4). The between-years comparison showed that growth resistance was significantly lower in 2019 (*p*<0.001) and 2020 (*p*<0.001) than in 2018, while the growth resistance in 2020 did not significantly differ from 2019. The same was true if we looked at clades of angiosperms and gymnosperms separately, however, for the gymnosperms the between-years comparison showed in addition a significant difference between 2019 and 2020 with growth resistance being lower in 2020 (*p*=0.031; Figure 4).

**Figure 4:**
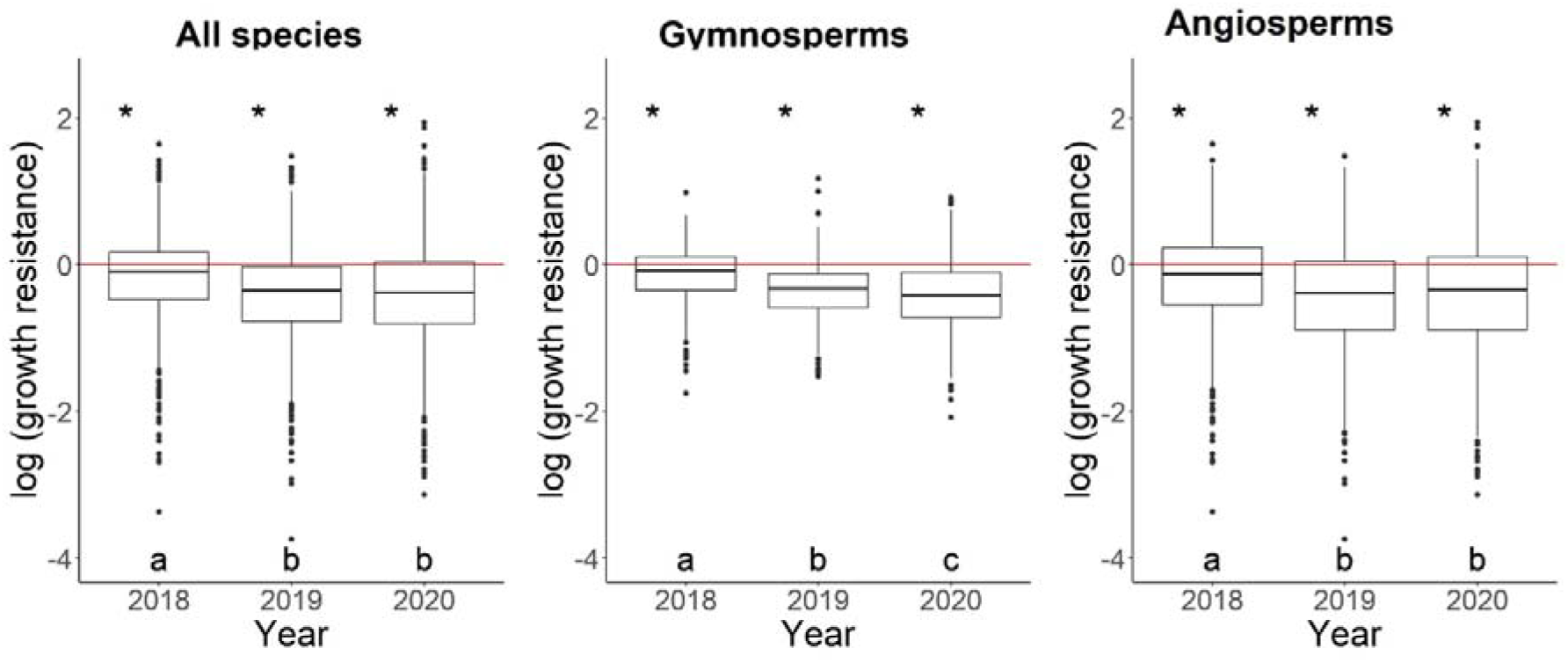
Growth resistance. Boxplots show the growth resistance of trees during the drought years 2018–2020 compared to the growth in the reference years (mean of 2016 and 2017) shown as red zero-line. Across species a significant (*p*<0.05) reduction in growth resistance, indicated with the asterisks, compared to the growth in the reference years was observed. The significant differences between the years were tested with a post-hoc test and are indicated by the characters (a, b, c). Similarly, significant (*p*<0.05) reductions in growth resistance were found when analysing the gymnosperms and the angiosperms separated.

### Functional trait responses

Looking at the effects of the single traits on the growth resistance, we found significant evidence for relationships within the gymnosperms as well as in the angiosperms (Figure 5, Table 1). Within the gymnosperms, we found that P50 had a significant negative effect on the growth resistance in 2019 (*p*<0.005; Figure 5, Table 1). Stomatal conductance had a significant negative effect on the growth resistance of gymnosperms in all three years (*p*=0.004, <0.001, =0.008, respectively). Also SLA, LNC and A_max_ had significant negative effects on the growth resistance of gymnosperms in all three years (*p*<0.001, for all), while the C:N significantly increased growth resistance in all three years (*p*=0.007, 0.029, <0.001, respectively). These models explained between 17–27 % of variation in growth resistance, through their fixed effects (marginal R^2^ (R^2^m)) and 42–48 % through their fixed and random effects (conditional R^2^ (R^2^c), Table 1).

**Figure 5:**
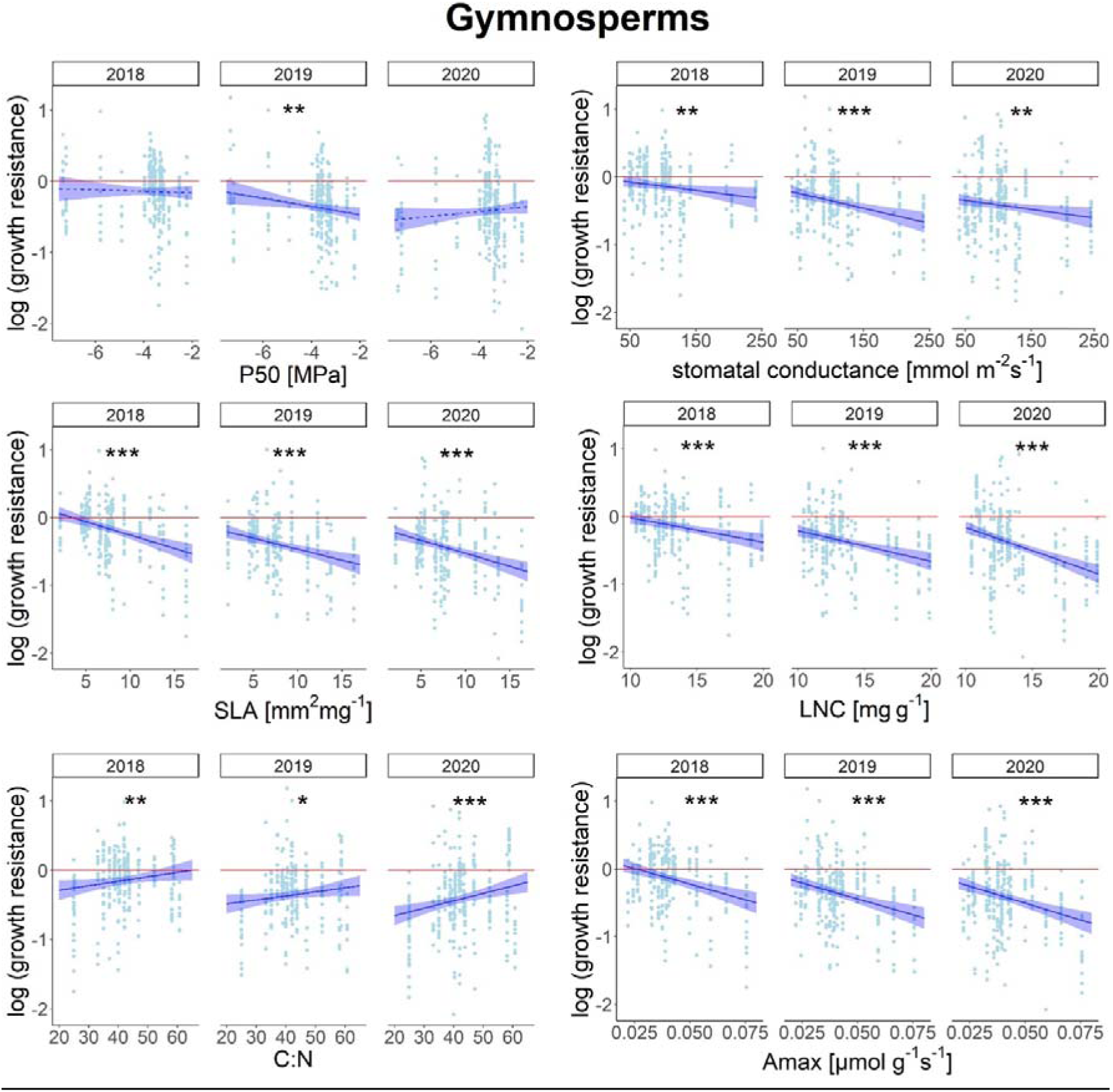
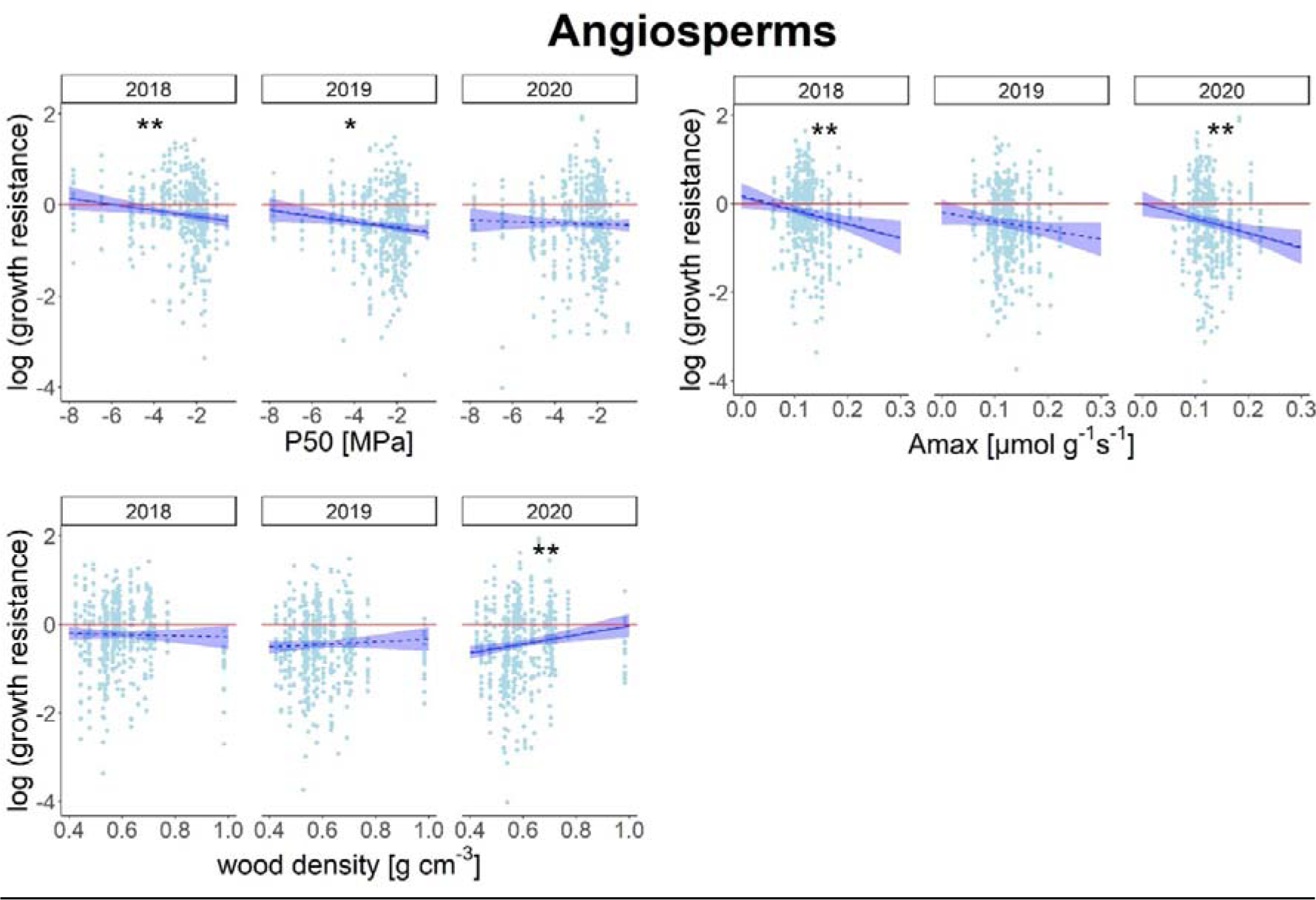
Trait-driven responses in growth resistance for gymnosperms and angiosperms. Shown are relationships between functional traits and the growth resistance of gymnosperm and angiosperm trees during the drought years 2018–2020 based on linear mixed-effects model fits. Growth resistance is depicted compared to tree growth in the reference years (mean of 2016 and 2017) shown as red zero-line. The asterisks indicate significant relationships (* *p*<0.05, ** *p*<0.01, *** *p*<0.001), while a dashed line symbolizes a non-significant relationship. Shaded bands show a 95% confidence interval.

**Table 1:** Growth resistance. explained by the single traits and the principal component 1 (PC1) for the drought years 2018–2020 for gymnosperms and angiosperms. Species number is the number of species include in the model for the single trait. Growth resistance is the slope of the relationship, with green and red indicating positive and negative relationships based on linear-mixed effects model fits, respectively. The marginal R^2^ (*R*^2^*m) shows the variation explained by fixed and the conditional R*^2^ (*R*^2^*c*) the variation explained by fixed and random effects. The asterisks indicate significant relationships (* *p*<0.05, ** *p*<0.01, *** *p*<0.001).

|  | Gymnosperms |  |  |  |  |  |  |  |  |
| --- | --- | --- | --- | --- | --- | --- | --- | --- | --- |
|  | spec no. | R<sup>2</sup>m | R<sup>2</sup>c | growth resistance 2018 | sig. | growth resistance 2019 | sig. | growth resistance 2020 | sig. |
| P50 | 23 | 0.18 | 0.43 | -0.010 |  | -0.057 | ** | 0.034 |  |
| stomata density | 23 | 0.17 | 0.42 | <0.001 |  | <-0.001 |  | 0.001 |  |
| stomatal conductance | 23 | 0.20 | 0.43 | -0.001 | ** | -0.002 | *** | -0.001 | ** |
| SLA | 21 | 0.27 | 0.48 | -0.040 | *** | -0.033 | *** | -0.039 | *** |
| LNC | 22 | 0.25 | 0.48 | -0.036 | *** | -0.045 | *** | -0.069 | *** |
| C:N | 23 | 0.19 | 0.43 | 0.006 | ** | 0.006 | * | 0.011 | *** |
| A<sub>max</sub> | 23 | 0.23 | 0.44 | -9.227 | *** | -9.424 | *** | -9.998 | *** |
| wood density | 23 | 0.17 | 0.42 | -0.030 |  | -0.030 |  | -0.572 |  |
| PC1 | 19 | 0.24 | 0.47 | 0.066 | *** | 0.059 | *** | 0.071 | *** |
|  | Angiosperms |  |  |  |  |  |  |  |  |
|  | spec no. | R<sup>2</sup>m | R<sup>2</sup>c | growth resistance 2018 | sig. | growth resistance 2019 | sig. | growth resistance 2020 | sig. |
| P50 | 43 | 0.08 | 0.29 | -0.065 | ** | -0.064 | * | -0.015 |  |
| stomata density | 45 | 0.07 | 0.28 | <-0.001 |  | <0.001 |  | <0.001 |  |
| stomatal conductance | 45 | 0.07 | 0.29 | <-0.001 |  | <-0.001 |  | <0.001 |  |
| SLA | 46 | 0.27 | 0.48 | -0.003 |  | -0.008 |  | -0.003 |  |
| LNC | 44 | 0.07 | 0.29 | 0.004 |  | 0.001 |  | -0.010 |  |
| C:N | 45 | 0.07 | 0.28 | 0.002 |  | 0.001 |  | 0.007 |  |
| A<sub>max</sub> | 46 | 0.08 | 0.29 | -3.166 | ** | -2.023 |  | -3.258 | ** |
| wood density | 46 | 0.08 | 0.30 | -0.142 |  | 0.297 |  | 1.013 | ** |
| PC1 | 39 | 0.07 | 0.29 | -0.048 |  | -0.047 |  | -0.051 |  |

For the angiosperms we found that P50 had a negative effect on the growth resistance in the year 2018 (*p*=0.005) and in 2019 (*p*=0.011; Figure 5, Table 1). A_max_ had a negative effect on the growth resistance for angiosperms in 2018 (*p*=0.003) and in 2020 (*p*=0.005) and wood density positively affected growth resistance in 2020 (*p*=0.003). These models explained between 7-27 % of variation in growth resistance through their fixed (R^2^m) and 28-48 % through their fixed and random effects (R^2^c, Table 1).

### Trait spaces

The principal component analysis (PCA) of all species within the trait space showed a clear separation between gymnosperms and angiosperms (Figure S3). Key drivers are the traits: SLA, LNC, C:N and A_max_ that clearly separated the two clades. Due to the strong separation in the trait space between the clades, we ran separate PCAs for both clades. When looking at the PCA for the gymnosperms only, we found that LES traits from the fast-slow-gradient are mainly associated with the first PCA axis (38 %; Figure 6), such as SLA, LNC, C:N, and A_max_. The first principal component (PC1) for gymnosperms as predictor, showed significant positive effects on the growth resistance for all three drought years (*p*<0.001, respectively; R^2^m = 24 %, R^2^c = 47%), meaning that gymnosperms with conservative traits featured a higher growth resistance (Figure 7, Table 1), which is in line with the single trait responses. The PCA of the angiosperms, also showed LES traits (SLA, LNC and C:N) mainly associated with the first PCA axis (31.54 %; Figure 6), but the PC1 as predictor did not significantly affect growth resistance (Figure 7, Table 1).

**Figure 6:**
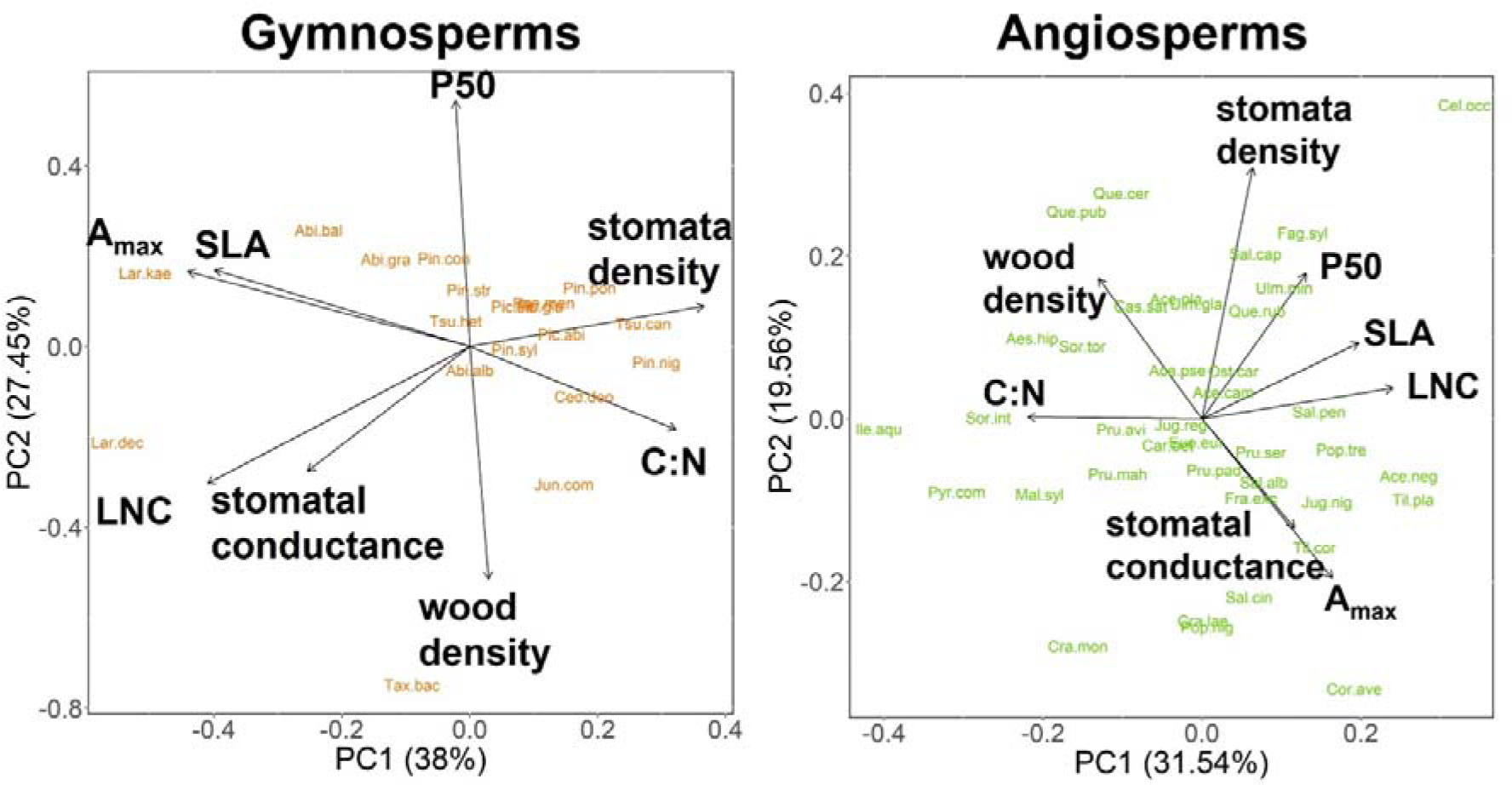
PCAs of gymnosperms and angiosperms depicting the trait space of the continuous traits P50, stomatal density, stomatal conductance, SLA, LNC, C:N, A_max_, and wood density.

**Figure 7:**
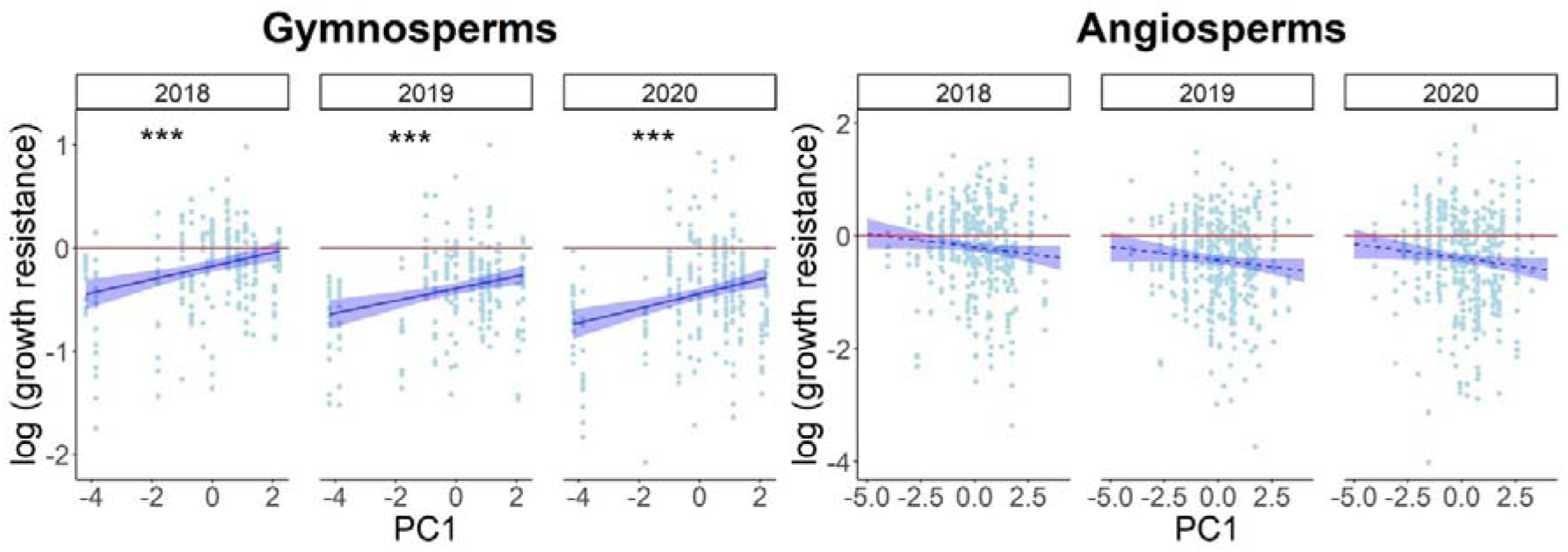
Principal components as predictors of tree growth resistance for gymnosperms and angiosperms. Shown are relationships between the principal component 1 (PC1) and growth resistance of gymnosperm and angiosperm trees during the drought years 2018–2020 based on linear mixed-effects model fits. Growth resistance is depicted compared to tree growth in the reference years (mean of 2016 and 2017) shown as red zero-line. The asterisks indicate significant relationships (* *p*<0.05, ** *p*<0.01, *** *p*<0.001), while a dashed line symbolizes non-significant relationships. Shaded bands show a 95% confidence interval.

For the gymnosperms, the traits P50 and wood density formed a gradient in opposing directions, thus higher P50 was associated with lower wood density. For the angiosperms, the two-dimensional trait space showed that P50 and wood density both load on PC2 in positive direction. However, when including PC4, P50 and wood density clearly showed an opposing pattern (Figure S4), that we also can see in the single trait responses (Figure 5). Stomata density and stomatal conductance also point, in both clades, in opposing direction and form a gradient of higher stomata density with lower stomatal conductance, which is for the gymnosperms on PC1 and for the angiosperms for PC2.

### Response types

With the single species models, we found recurring patterns that allowed us to classify the species *a posteriori* based on their drought responses over the three consecutive drought years into four main response classes: ‘Sufferer’, ‘Late sufferer’, ‘Resister’ and ‘Recoverer’ (Figure 8, Figure S5). As ‘Sufferers’ we defined species with a significant negative growth resistance in all three years 2018, 2019, and 2020. ‘Late sufferers’ are species that had no significantly reduced growth resistance initially in 2018 but then a significantly reduced growth resistance latest in 2020. Species defined as ‘Resisters’ had no significantly reduced growth resistance in 2018, 2019, and 2020. The ‘Recoverers’ are species that had significantly negative growth resistance in 2018 or/and 2019; and had no significantly reduced growth resistance in 2020 (Figure 8). The full decision tree behind this classification is shown in Figure S5, while the classification for every single species is listed in Table S1.

**Figure 8:**
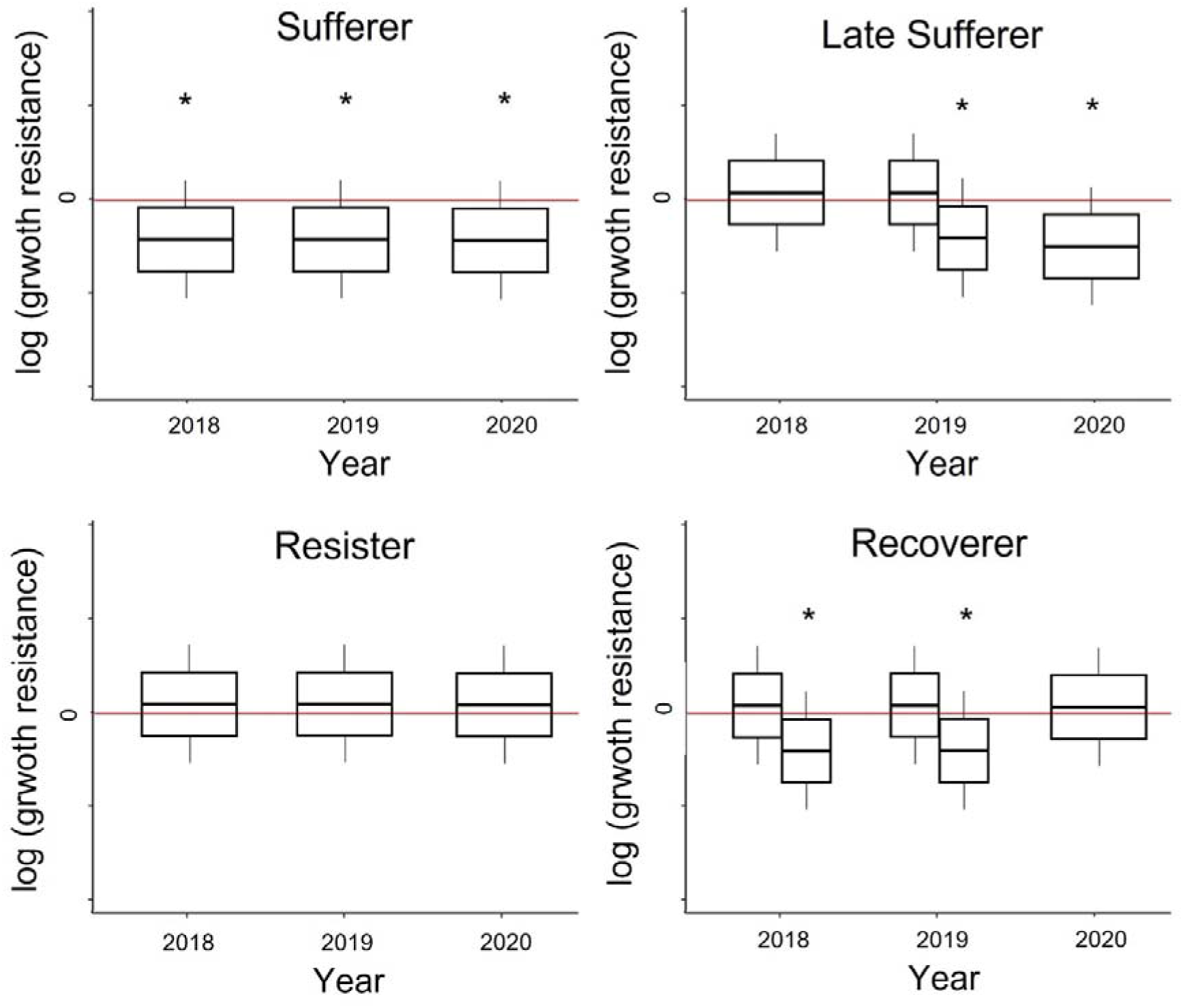
Response type classification. Growth response patterns over the three drought years 2018–2020 for the four response types classified due to the decision tree in Figure S4. The boxplots above the red reference zero line represent positive or not significant resistance and the ones below the red zero line negative resistance values with asterisks indicating a significant response in growth resistance. The divided boxplot for the Late Sufferer in 2019 and for the Recoverer in 2018 and 2019 show positive and not significant (1^st^ boxplot) or significant negative (2^nd^ boxplot) effects, since they represent two divergent paths of the decision tree (Figure S5). In addition, for the Recoverer, either 2018 or 2019 or both years needed to be significantly negative as shown in the decision tree (Figure S5).

We observed clear patterns of how these response types are distributed over the phylogenetic clades and that they are statistically independent from each other (Figure 9; *p*=0.149). The 23 gymnosperms did mainly show a growth pattern of ‘Sufferer’ (8 species) and ‘Late sufferer’ (10 species), and had only 2 species counting as ‘Recoverer’ and 3 ‘Resister’ species. Within the 48 angiosperms we found 11 ‘Sufferer’ and 13 ‘Late sufferer’, but also 13 ‘Recoverer’ and 11 ‘Resister’. However, we did not detect an apparent pattern of the four response types within the trait spaces of gymnosperms and angiosperms (Figure S6).

**Figure 9:**
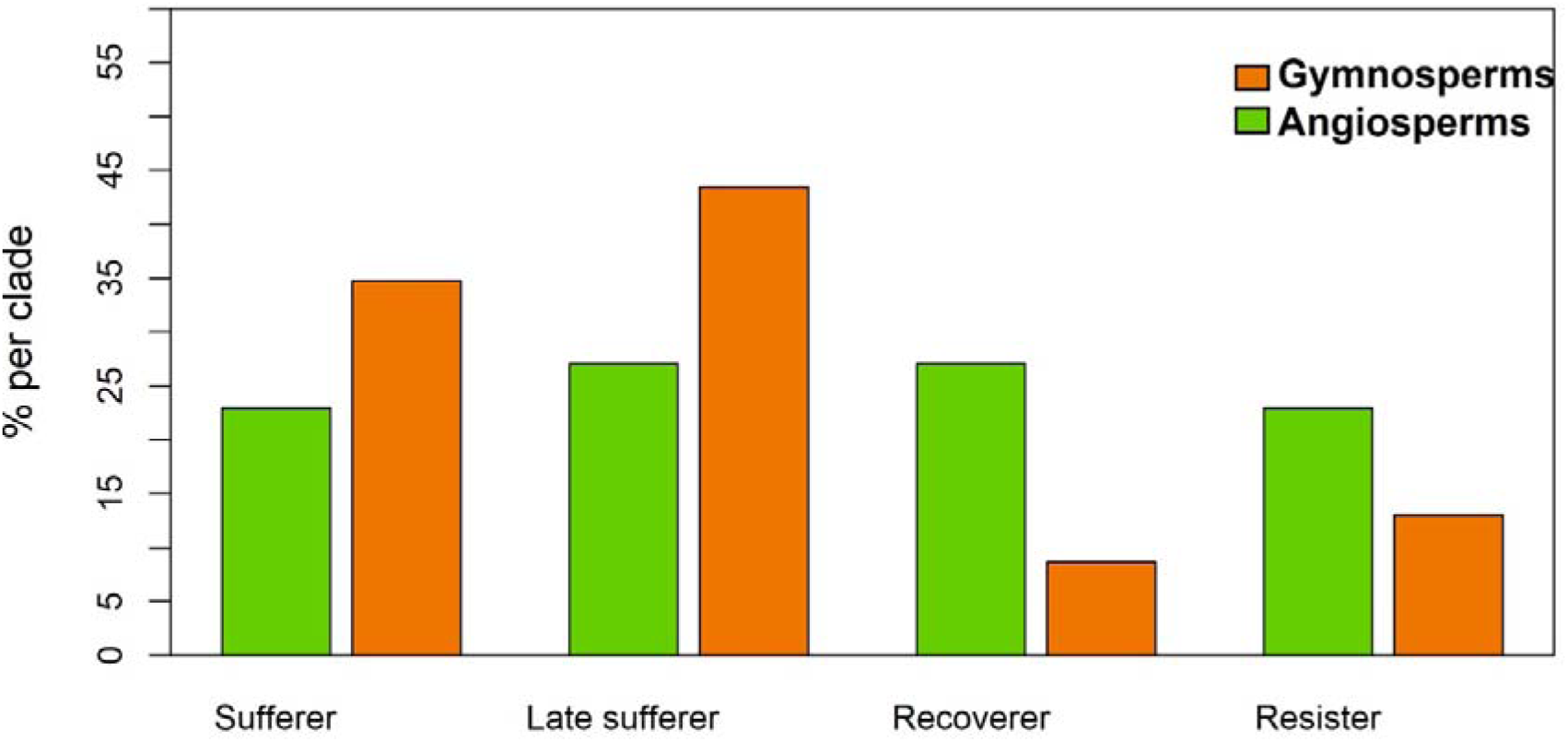
Distribution of clades into response types. ‘Sufferer’, ‘Late sufferer’, ‘Recoverer’, and ‘Resister’ separated for the clades of angiosperms and gymnosperms. Fisher’s exact test showed significant differences between clades and response types (*p*=0.149).

## Discussion

### Tree growth resistance across species

Over the three consecutive drought years 2018–2020, we found evidence for significant growth reductions in our analysis of 71 tree species growing at a single site under the controlled experimental conditions of the research arboretum ARBOfun. Overall, we confirmed our first hypothesis that the 2018–2020 drought caused a growth reduction over the whole drought period, but growth reductions were greater in 2019 and 2020. During drought, trees lack water and face the trade-off between carbon gain and water loss. Thus, growth reduction is a reaction to maintain physiological key processes which prevent the tree from carbon starvation and hydraulic failure, as the two major reasons for tree mortality during droughts (Adams et al., 2017; N. McDowell et al., 2008; Sala et al., 2010; Schuldt et al., 2020; Sevanto et al., 2014). In contrast to most previous studies (but see Liu et al. (2022)), the use of shoot increments as a growth indicator enabled us to precisely measure tree growth even though the trees at our study site are relatively young and thus not suitable for taking tree cores for standard dendrochronological analysis. Given that climatic drought severity was lower in the consecutive drought years 2019 and 2020 (Figure 2), the stronger growth reduction in 2019 and 2020 is likely the result of drought legacy effects. Especially the 23 species of the response type ’Late sufferer’ emphasize the progressive deterioration of growing conditions under consecutive drought due to legacy effects, since they did not show reduced growth during the first drought year (2018), while they had reduced growth in the later years (2019/2020; Table S1). Legacy effects, such as damages to the tree’s water transport system, are known to effect trees and forests negatively up to five years after the drought (Anderegg et al., 2013; Anderegg et al., 2015; Bigler et al., 2006; Gazol et al., 2020; Kannenberg et al., 2018; Schnabel et al., 2022). We also found that the growth resistance for the gymnosperms was more strongly reduced compared to the angiosperms in all three years (Figure 4). This is also supported by the response type classification where most gymnosperm species (>75 %), except for five species, suffered during the consecutive drought years and were therefore classified either as ‘Sufferer’ or as ‘Late sufferer’, while of the angiosperms only 50 % were classified as ‘Sufferer’ or ‘Late sufferer’ (Figure 9, Table S1). Already others found evidence that gymnosperms suffer more strongly during drought, since for gymnosperms reinvesting into damaged leaves is costly (Anderegg et al., 2020; Larysch et al., 2022; Song et al., 2022). However, also equally high mortality risks during drought were found for angiosperms and gymnosperms worldwide (Anderegg et al., 2016).

### Drought-tolerance traits and wood density

For the drought tolerance trait P50, which we expected to be negatively related to growth resistance, we found a significantly negative effect on growth resistance of gymnosperms in 2019 and of angiosperms in 2018 and 2019 (Figure 5). Thus, we could confirm our second hypothesis as we found that species whose functional traits indicate increasing drought tolerance by the P50 trait show an increase in embolism resistance and, hence, also growth resistance (Guillemot et al., 2022; Petruzzellis et al., 2022).

For angiosperms, increasing wood density increased growth resistance in 2020, as we expected, but we did not observe this relationship for gymnosperms, which might be due to a much smaller range of wood densities within the gymnosperms (0.40-0.65 g cm^−3^) compared to angiosperms (0.43-0.98 g cm^−3^). For gymnosperms, wood density was negatively related to P50 (PC2, Fig. 6), similarly a strong negative correlation between wood density and P50 existed for angiosperms (albeit on PC4, Table S2). Thus, a low P50, meaning high embolism resistance, links to high wood density, causing increased growth resistance. This supports previous evidence, that wood density is associated with other drought tolerance traits such as hydraulic safety margin and P50 (Oliveira et al., 2021; Rosner, 2017). In addition, we found that wood density and P50 formed a separate axis independent to the leaf economics spectrum (LES, Díaz et al., 2016). Although the overall effect of wood density on growth resistance is still debated (Chave et al., 2009; L. Poorter, 2008), our study provides evidence that wood density is associated with enhanced growth resistance.

For gymnosperms and angiosperms, stomata density and stomatal conductance loaded in opposing directions in the PCAs (Figure 6), which implies that species with low stomata density have high stomatal conductance, most likely due to few but large stomata. As expected, gymnosperms with lower stomatal conductance had a higher growth resistance. Moreover, for the gymnosperms, effects of stomatal conductance on growth resistance were similar as for the LES traits (SLA, LNC, C:N, A_max_), which is likely related to stomata density and stomatal conductance being associated with these LES traits in trait space (PC1, Fig. 6), an association which has been reported previously albeit for angiosperms in the subtropics (Kröber et al., 2014). Thus, we expect a high stomatal conductance to be associated with acquisitive resource use. However, for angiosperms, we did not find such close association between stomata traits and LES traits nor with hydraulic traits (P50).

### Leaf economics spectrum traits

In gymnosperms, LES trait expressions associated with conservative resource use and slower growth increased growth resistance during the drought (Figure 5, Table 1). In addition, these LES traits formed an important axis of functional variation on the first axis (i.e., SLA, LNC, C:N, A_max_), ranging from fast to slow strategies. As expected, PC1 was positively related to growth resistance within all three years (Figure 7, Table 1). For angiosperms, we found significantly negative growth responses for A_max_ in 2018 and 2020 (Figure 5, Table 1), showing that species with a high light-saturated maximum photosynthetic rate - usually associated with fast growth - have a low growth resistance. Also, for angiosperms, most of the LES traits loaded on the first PCA axis (i.e., SLA, LNC, and C:N, Figure 6). Thus, we confirmed hypothesis 3 for gymnosperms and angiosperms (albeit weaker associations of LES traits were observed). This means species with resource acquisition traits favouring rapid growth are more susceptible to drought and show a stronger reduction in growth resistance during consecutive drought years. Earlier studies suggested that LES trait expressions related to conservative resource use and slow growth are related to (1) a lower drought mortality across biomes (Greenwood et al., 2017), and (2) a higher drought tolerance in the tropics (Guillemot et al., 2022). Similarly to our findings, a study in subtropical experimental tree communities reported recently that acquisitive species had reduced growth resistance under drought conditions (Schnabel et al., 2024), albeit based on fewer tree species. The weaker trend for the angiosperms in our study could be caused by the fact that 50 % of the angiosperms did not suffer substantially during the entire drought period. Thus, according to our response type classification, 27 % of the angiosperms recover already during the drought, while 23 % resist the drought in their growth response (Figure 9, Table S1). Further, our study shows that even though we found a legacy effect in the growth resistance in 2019 and 2020, the LES trait control was directly present from the first drought year of 2018 onwards. For the first time, we report clear evidence for LES traits driving tree growth resistance for a wide species set under nearly identical growing conditions and extreme drought conditions, causing legacy effects.

### Management

Trees in the arboretum ARBOfun were planted at a wide spacing, which prevented tree-tree interactions. Thus, our results can be interpreted as the intrinsic, trait-driven response of the species to climatic conditions without influences of competition, competitive reduction or facilitation (Forrester & Pretzsch, 2015), which are otherwise present in forests and shape effects of functional traits on ecosystem functioning (Trogisch et al., 2021). Our study thus captures ‘pure’ trait-driven responses of a wide set of Central European tree species to consecutive drought years including species dominating today’s forests but also those which may dominate under a future climate regime such as currently subordinate or biographically neighbouring tree species. The traits and trait syndromes (such as P50 and the LES) we observed to influence growth resistance can thus inform management decisions on tree species choice and be used to improve the predictive capacity of forest models. The identification of the four response types helps to recognize growth resistance pattern across species, but also gives important insights for single key species. The two currently economically most relevant tree species in Central European managed forest, *P. abies* and *P. sylvestris,* together making up, for instance, 47.7 % of Germany’s managed forest (BWI 2012), suffered strongly the last years (Senf et al., 2020). They showed a drought response of ‘Late sufferer’, which indicates that they likely strongly suffer in the coming century facing more regular and more intense droughts caused by climate change (IPCC, 2014). Similar negative predictions were also found by others (Buras & Menzel, 2019; Kölling & Mette, 2022; Wessely et al., 2024). In contrast, the angiosperms *F. sylvatica* and *Q. robur*, currently accounting for 25.8% of Germany’s managed forests (BWI 2012), showed response types of ‘Recoverer’ and ‘Resister’, respectively. Also Kölling & Mette (2022) and Buras & Menzel (2019) classified those two species as more resistance against climate change. While the drought resistance of *F. sylvatica* is under debate (Kunz et al., 2018), we could reinforce evidence for it. Two species of currently minor merchantable value, but with potential to gain in economic importance for Central European forests in the future are *S. torminalis* and *Q. pubescens* (Buras & Menzel, 2019; Kunz et al., 2018). We also classified those as the response type of ‘Recoverer’ and ‘Resister’, respectively. Thus, our response type classification approach, helps to depict single species responses, even though we did not find clear patterns of the response types within the trait spaces (Figure S6), pointing to the fact that similar responses may be achieved by different but equivalent trait configurations which warrants further investigation.

### Reflection

We did not explicitly correct for phylogeny, since the separation of clades (angiosperms and gymnosperms) already captures a large portion of the phylogenetic signal (see Figure S3). Further, we also did not control for tree size. One would expect larger fast-growing trees to root deeper and thus have better water access, however we found fast-growing species are less growth resistant. The gap-filling of the trait data from the TRY database is a helpful and indispensable tool to be able to investigate many traits for a wide set of species. However, it has the weakness that traits for different species in TRY have been measured with different methods, at different times and places, which can, dependent on the species and the trait, induce a high amount of variation due to strong plasticity. Moreover, particularly P50, which is a key trait for drought tolerance (Choat et al., 2012), is difficult to measure, especially in ring-porous species with very long vessels. Therefore, we excluded P50 values larger than −0.5, as suggested by (Sergent et al., 2020), due to unrealistically high values. However, we decided against excluding P50 values smaller than leaf turgor loss point (P_tlp_) values, as suggested by (Guillemot et al., 2022), due to the fact, that the available trait data on those two traits in the TRY database came mainly from different studies and did not cover all our species. Overall, our trait-based models explained only moderate shares of variation in growth resistance with a higher predictive capacity for gymnosperms compared to angiosperms (Table 1), but we expect that with more and particularly in-situ measured traits such models are likely to increase in their predictive capacity. Similarly, with such an enhanced trait coverage, we might eventually be able to derive trait-based predictions for the assignment of species to the observed response types.

## Outlook and conclusion

For future studies, we plan for in-situ functional trait measurements which likely have the potential to improve growth predictions under consecutive hotter droughts. Moreover, besides the drought tolerance and LES traits we studied here, other hydraulic traits such as turgor loss point or hydraulic safety margin, but especially also belowground traits may be important predictors of growth resistance to drought. Belowground traits such as specific root length, root tissue density or root C:N, which capture a conservation, a collaboration and a plant size gradient (Bergmann et al., 2020; Comas et al., 2013; Weigelt et al., 2021) were already found to affect above- and belowground plant productivity under drought (Brunner et al., 2015; Comas et al., 2013). Thus, future studies should consider more and especially belowground traits. The importance of drought tolerance and LES traits for growth resistance, and the recovery of some species under consecutive drought as we have shown, suggest that functional traits might also explain growth resilience. Some species, such as *F. sylvatica*, *Quercus rubra* or *S. torminalis* did already recover during the drought, even though an overarching legacy effect was visible. However, we do not know whether and when the species of ‘Sufferers’, such as *Larix decidua* or *Ulmus laevis* and the ‘Late sufferers’, such as *Acer campestre*, *P. abies* or *P. sylvestris* do recover over time. Hence, studying growth resilience and recovery including also the recent wetter years 2023 or even 2024 is hence of high interest at our study site. Overall, we are planning on future studies looking at tree growth expression over pre-drought, consecutive hotter drought and post-drought years, studying resistance, recovery and resilience to these contrasting climatic conditions and to further explore the underlying, trait-based mechanisms by including in-situ measured trait data capturing both, above- and belowground trait gradients.

In conclusion, we observed significantly reduced growth across the 71 tree species during the consecutive hotter drought years 2018–2020 with legacy effects further reducing growth resistance during 2019 and 2020. Drought-tolerance and LES traits were important predictors of growth resistance, with lower growth resistance observed in species featuring trait expressions indicative of low drought tolerance (high P50), fast growth and acquisitive resource use (high SLA, LNC, and A_max_). Trait-growth resistance relationships were clearer for gymnosperms than for angiosperms. We expect these findings to facilitate the development of management strategies for forests under a future climate regime characterized by more frequent, severe and prolonged droughts through supporting tree species choice and the improvement of forest models.

## Supporting information

SupportingInformation

## Acknowledgement

We thank the TRY database for the provision of trait data.

## Author Contributions

**LK** formal analysis, visualization, writing – original draft preparation; **FS** conceptualization, formal analysis, methodology, supervision, writing – review & editing; **RR** conceptualization, formal analysis, writing – review & editing; **AR** investigation, writing – review & editing; **JK** methodology, formal analysis, writing – review & editing; **KA** formal analysis, writing – review & editing; **AK** investigation, writing – review & editing; **TK** investigation, writing – review & editing; **CW** idea, conceptualization, supervision, funding acquisition, project administration, writing – review & editing

## Date Availability Statement

We plan to publish the data at iBID (iDiv).

