## SupportingInformation for "Functional traits explain growth resistance to successive hotter droughts across a wide set of common and future tree species in Europe"

**Supporting Information**

Figure S1

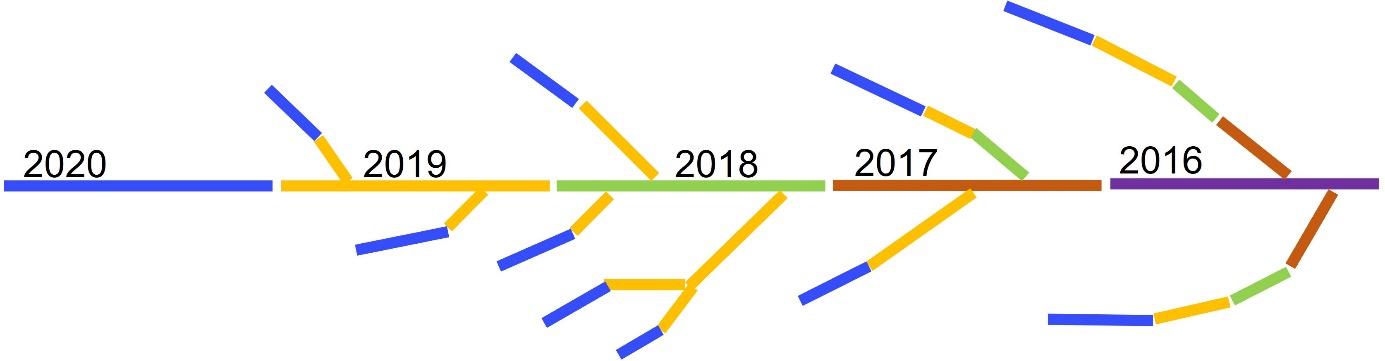

**Figure S1:** Sketch of a branch showing the shoot increment per year back to 2016. Shoot increment was always measured at the main branch.

Figure S2

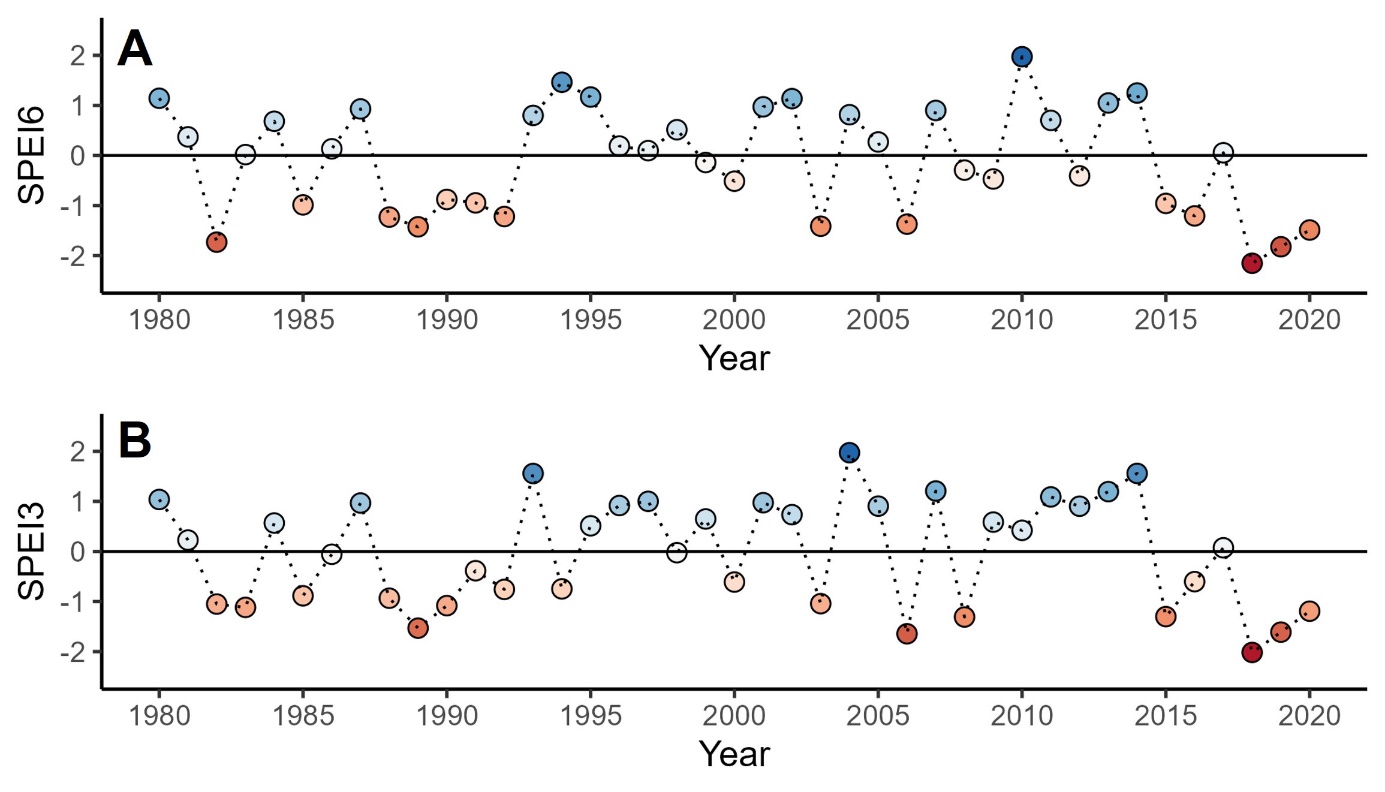

**Figure S2**: Standardised Precipitation-Evapotranspiration Index (SPEI) calculated for three (SPEI3) and six months (SPEI6). The SPEI6 was calculated for the six months of the vegetation period (April to September) for each year. The SPEI3 was calculated for the three months of the peak vegetation period (May to July) for each year. The zero line is the reference period 1981–2010. Blue coloured dots indicate positive SPEI values, while red coloured dots show negative SPEI values.

Figure S3
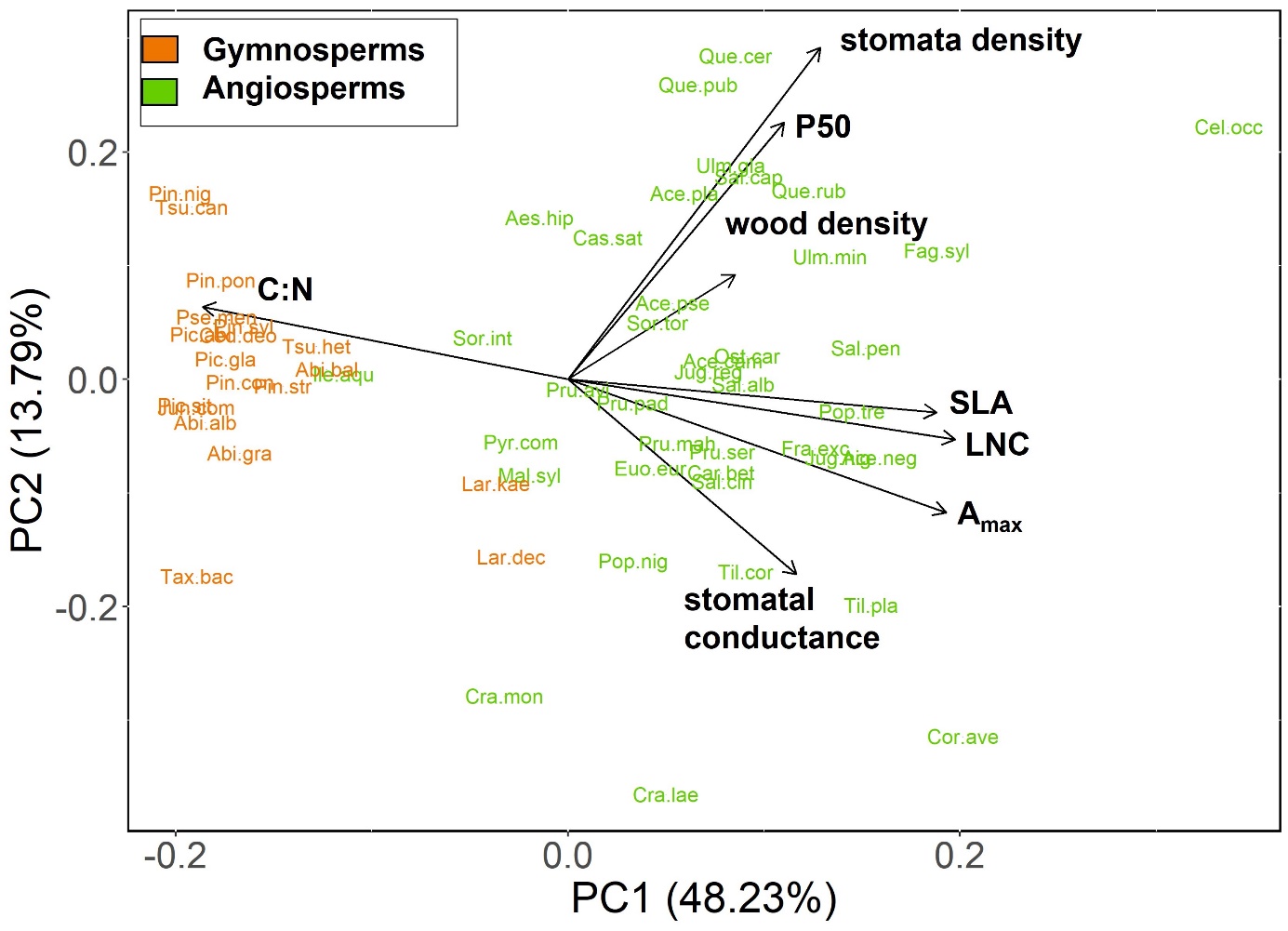

**Figure S3**: PCA of all species depicting the trait space of the continuous variables SLA, LNC, C:N, A_max_, stomata density, stomatal conductance, P50 and wood density.

Figure S4

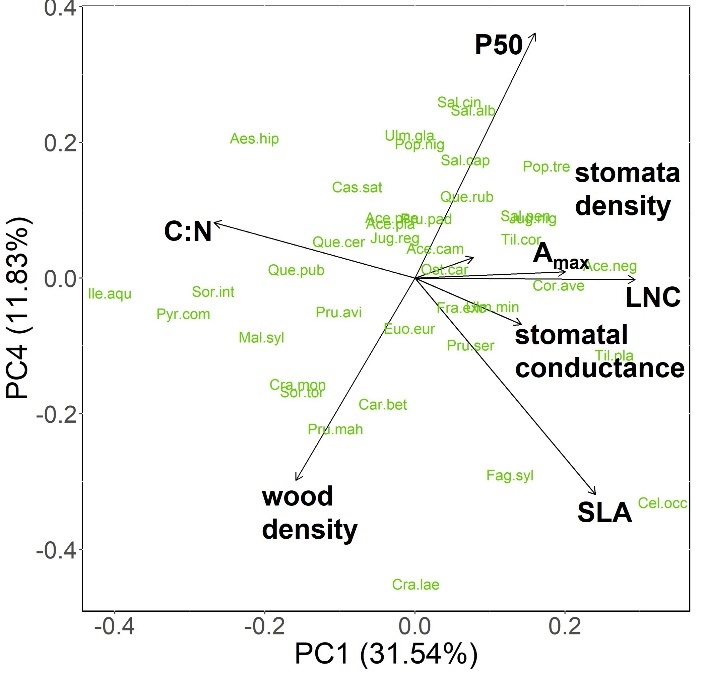

**Figure S4**: PCA of angiosperms (PC1 and PC4) depicting the trait space of the continuous traits P50, stomatal density, stomatal conductance, SLA, LNC, C:N, A_max_, and wood density.

Figure S5
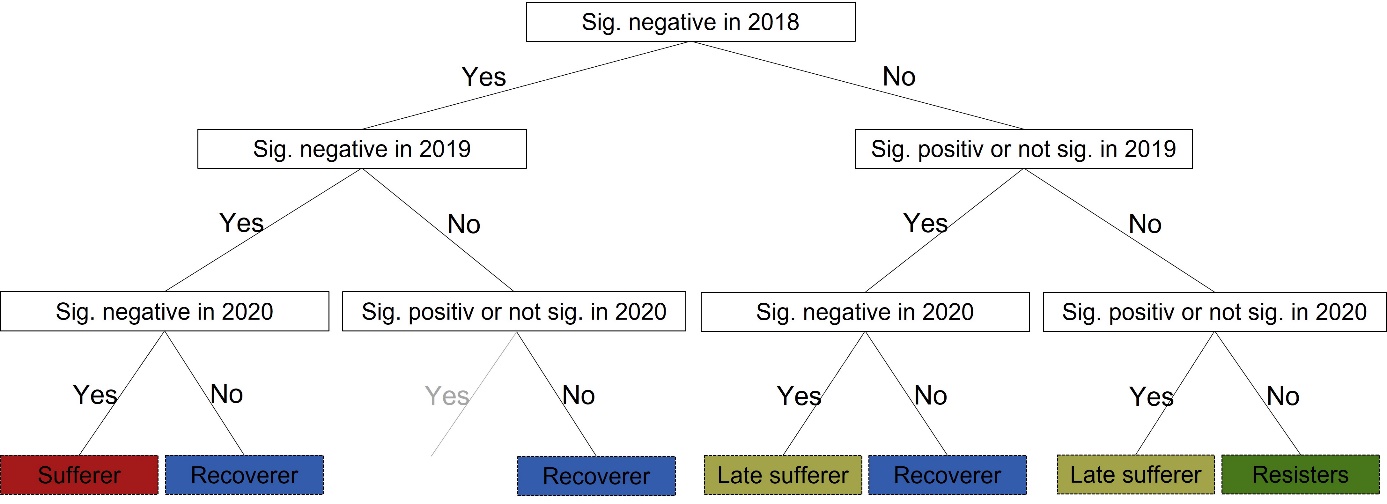

**Figure S5**: Decision tree for the classification of the response types, based on the linear mixed-effect models of the single species. The final classification due to the four response types ‘Sufferer’, ‘Late sufferer’, ‘Recoverer’ and ‘Resisters’ shows the reaction patterns of growth responses during the three drought years. The grey line did not occure for any of the investigated species.

Figure S6
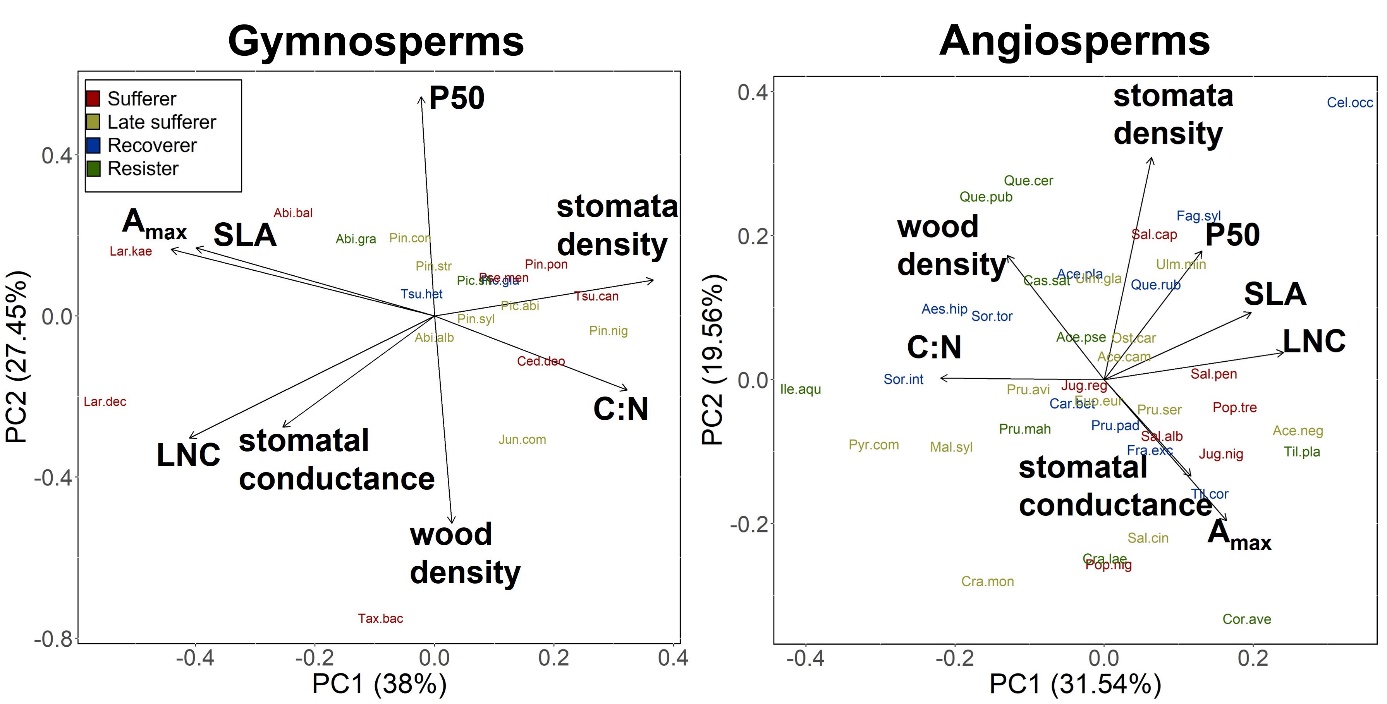
 **Figure S6**: PCAs of gymnosperms and angiosperms depicting the trait space of the continuous variables SLA, LNC, C:N, A_max_, stomata density, stomatal conductance, P50 and wood density. Colour due to the response types.

**Table S1**: Species list of all investigated species and their attribution to the response types. The green colours indicate not significant and significant positive growth resistance in the years 2018, 2019 and 2020, while the red colour indicate significant negative growth resistance. The response types result from the growth resistance pattern according to the decision tree in Figure S5. R^2^m and R^2^c are the predictive power of the single species model, without and with the random effect included, respectively.

| **Species** | **Response type** | Growth resistance 2018 | Growth resistance 2019 | Growth resistance 2020 | R^2^m | R^2^c |
| --- | --- | --- | --- | --- | --- | --- |
| *Abies alba* Mill. | Late sufferer |  |  |  | 0.37 | 0.48 |
| *Abies balsamea* (L.) Mill. | Sufferer |  |  |  | 0.44 | 0.59 |
| *Abies grandis* (Douglas ex D. Don) Lindley | Resister |  |  |  | 0.03 | 0.49 |
| *Acer campestre* L. | Late sufferer |  |  |  | 0.19 | 0.44 |
| *Acer negundo* L. | Late sufferer |  |  |  | 0.31 | 0.45 |
| *Acer platanoides* L. | Recoverer |  |  |  | 0.33 | 0.50 |
| *Acer pseudoplatanus* L. | Resister |  |  |  | 0.02 | 0.41 |
| *Acer saccharinum* L. | Sufferer |  |  |  | 0.32 | 0.42 |
| *Aesculus hippocastanum* L. | Recoverer |  |  |  | 0.17 | 0.38 |
| *Carpinus betulus* L. | Recoverer |  |  |  | 0.13 | 0.25 |
| *Castanea sativa* Mill. | Resister |  |  |  | 0.03 | 0.03 |
| *Cedrus deodara* (Roxb. ex D.Don) G.Don | Sufferer |  |  |  | 0.37 | 0.41 |
| *Cedrus libani* A. Rich. | Late Sufferer |  |  |  | 0.36 | 0.36 |
| *Celtis occidentalis* L. | Recoverer |  |  |  | 0.25 | 0.42 |
| *Corylus avellana* L. | Resister |  |  |  | 0.08 | 0.20 |
| *Corylus colurna* L. | Recoverer |  |  |  | 0.33 | 0.62 |
| *Crataegus laevigata* (Poir.) DC. | Resister |  |  |  | 0.06 | 0.32 |
| *Crataegus monogyna* Jacq. | Late sufferer |  |  |  | 0.11 | 0.22 |
| *Euonymus europaeus* L. | Late sufferer |  |  |  | 0.21 | 0.46 |
| *Fagus sylvatica* L. | Recoverer |  |  |  | 0.09 | 0.27 |
| *Fraxinus excelsior* L. | Recoverer |  |  |  | 0.08 | 0.52 |
| *Ilex aquifolium* L. | Resister |  |  |  | 0.09 | 0.09 |
| *Juglans nigra* L. | Sufferer |  |  |  | 0.37 | 0.37 |
| *Juglans regia* L. | Sufferer |  |  |  | 0.41 | 0.60 |
| *Juniperus communis* L. | Late sufferer |  |  |  | 0.13 | 0.39 |
| *Larix decidua* Mill. | Sufferer |  |  |  | 0.54 | 0.65 |
| *Larix kaempferi* (Lamb.) Carr. | Sufferer |  |  |  | 0.50 | 0.69 |
| *Malus sylvestris* (L.) Mill | Late sufferer |  |  |  | 0.16 | 0.50 |
| *Mespilus germanica* L. | Recoverer |  |  |  | 0.33 | 0.58 |
| *Ostrya carpinifolia* Scop. | Late sufferer |  |  |  | 0.24 | 0.34 |
| *Picea abies* (L.) H. Karst. | Late sufferer |  |  |  | 0.22 | 0.22 |
| *Picea glauca* (Moench) Voss | Recoverer |  |  |  | 0.46 | 0.52 |
| *Picea pungens* Engelm. | Late sufferer |  |  |  | 0.38 | 0.49 |
| *Picea sitchensis* (Bong.) Carr. | Resister |  |  |  | 0.14 | 0.57 |
| *Pinus cembra* L. | Late sufferer |  |  |  | 0.48 | 0.58 |
| *Pinus contorta* Douglas | Late sufferer |  |  |  | 0.09 | 0.54 |
| *Pinus mugo* Turra | Resister |  |  |  | 0.28 | 0.33 |
| *Pinus nigra* J.F. Arnold | Late sufferer |  |  |  | 0.49 | 0.70 |
| *Pinus ponderosa* Douglas ex C. Lawson | Sufferer |  |  |  | 0.47 | 0.53 |
| *Pinus strobus* L. | Late sufferer |  |  |  | 0.10 | 0.45 |
| *Pinus sylvestris* L. | Late sufferer |  |  |  | 0.44 | 0.44 |
| *Platanus acerifolia* (Aiton) Willd. | Sufferer |  |  |  | 0.38 | 0.47 |
| *Populus nigra* L. | Sufferer |  |  |  | 0.48 | 0.50 |
| *Populus tremula* L. | Sufferer |  |  |  | 0.20 | 0.25 |
| *Prunus avium* L. | Late sufferer |  |  |  | 0.15 | 0.36 |
| *Prunus mahaleb* L. | Resister |  |  |  | 0.07 | 0.12 |
| *Prunus padus* L. | Recoverer |  |  |  | 0.15 | 0.41 |
| *Prunus serotina* Ehrh. | Late sufferer |  |  |  | 0.35 | 0.35 |
| *Pseudotsuga menziesii* (Mirbel) Franco | Sufferer |  |  |  | 0.63 | 0.78 |
| *Pyrus communis* L. | Late sufferer |  |  |  | 0.35 | 0.35 |
| *Quercus cerris* L. | Resister |  |  |  | 0.06 | 0.17 |
| *Quercus pubescens* Willd. | Resister |  |  |  | 0.09 | 0.17 |
| *Quercus robur* L. | Resister |  |  |  | 0.06 | 0.10 |
| *Quercus rubra* L. | Recoverer |  |  |  | 0.32 | 0.51 |
| *Robinia pseudoacacia* L. | Sufferer |  |  |  | 0.27 | 0.55 |
| *Salix alba* L. | Sufferer |  |  |  | 0.32 | 0.32 |
| *Salix caprea* L. | Sufferer |  |  |  | 0.32 | 0.32 |
| *Salix cinerea* L. | Late sufferer |  |  |  | 0.28 | 0.31 |
| *Salix pentandra* L. | Sufferer |  |  |  | 0.77 | 0.81 |
| *Sorbus domestica* L. | Resister |  |  |  | 0.14 | 0.38 |
| *Sorbus intermedia* (Ehrh.) Pers. | Recoverer |  |  |  | 0.15 | 0.26 |
| *Sorbus torminalis* (L.) Crantz | Recoverer |  |  |  | 0.30 | 0.55 |
| *Taxus baccata* L. | Sufferer |  |  |  | 0.32 | 0.41 |
| *Tilia cordata* Mill. | Recoverer |  |  |  | 0.17 | 0.31 |
| *Tilia platyphyllos* Scop. | Resister |  |  |  | 0.11 | 0.53 |
| *Tilia tomentosa* Moench | Late sufferer |  |  |  | 0.14 | 0.33 |
| *Tsuga canadensis* (L.) Carrière | Sufferer |  |  |  | 0.69 | 0.73 |
| *Tsuga heterophylla* (Raf.) Sarg. | Recoverer |  |  |  | 0.16 | 0.54 |
| *Ulmus glabra* Huds. | Late sufferer |  |  |  | 0.27 | 0.55 |
| *Ulmus laevis* Pall. | Sufferer |  |  |  | 0.26 | 0.34 |
| *Ulmus minor* Mill. | Late sufferer |  |  |  | 0.14 | 0.40 |

**Table S2**: PC loadings of the PCAs depicting the gymnosperms and angiosperms. Loadings for PC1, PC2, PC3 and PC4 are shown.

| Gymnosperms | | | | |
| --- | --- | --- | --- | --- |
|  | PC1 | PC2 | PC3 | PC4 |
| Standard deviation | 1.744 | 1.482 | 1.024 | 0.725 |
| Proportion of Variance | 0.380 | 0.275 | 0.131 | 0.066 |
| Cumulative Proportion | 0.380 | 0.655 | 0.786 | 0.851 |
|  | PC1 | PC2 | PC3 | PC4 |
| P50 | -0.025 | **0.598** | -0.149 | 0.088 |
| stomata density | **0.403** | 0.098 | **-0.455** | -0.207 |
| stomatal conductance | -0.280 | -0.304 | **-0.603** | **-0.470** |
| SLA | **-0.439** | 0.186 | -0.356 | 0.241 |
| LNC | **-0.451** | -0.333 | 0.066 | -0.186 |
| C:N | 0.355 | -0.202 | **-0.505** | **0.483** |
| A_max_ | **-0.485** | 0.182 | -0.127 | **0.433** |
| wood density | 0.032 | **-0.566** | 0.071 | **0.464** |
| Angiosperms | | | | |
|  | PC1 | PC2 | PC3 | PC4 |
| Standard deviation | 1.589 | 1.251 | 1.152 | 0.973 |
| Proportion of Variance | 0.315 | 0.196 | 0.166 | 0.118 |
| Cumulative Proportion | 0.315 | 0.511 | 0.677 | 0.795 |
|  | PC1 | PC2 | PC3 | PC4 |
| P50 | 0.277 | 0.378 | -0.249 | **0.625** |
| stomata density | 0.135 | **0.653** | -0.251 | 0.052 |
| stomatal conductance | 0.246 | -0.283 | **-0.640** | -0.119 |
| SLA | **0.417** | 0.198 | 0.091 | **-0.553** |
| LNC | **0.509** | 0.080 | 0.270 | -0.004 |
| C:N | **-0.464** | 0.005 | -0.277 | 0.142 |
| A_max_ | 0.347 | **-0.414** | **-0.416** | 0.015 |
| wood density | -0.275 | 0.365 | -0.367 | **-0.517** |
